# Thymidine kinase-independent click chemistry DNADetect™ probes for DNA proliferation assessment in malaria parasites

**DOI:** 10.1101/2023.03.26.534296

**Authors:** David H. Hilko, Gillian M. Fisher, Russell S. Addison, Katherine T. Andrews, Sally-Ann Poulsen

**Affiliations:** Griffith Institute for Drug Discovery, Griffith University, Nathan, Brisbane, Queensland 4111, Australia; School of Environment and Science, Griffith University, Nathan, Brisbane, Queensland 4111, Australia

## Abstract

Metabolic chemical probes are small molecule reagents that utilise naturally occurring biosynthetic enzymes for *in situ* incorporation into biomolecules of interest. These reagents can be used to label, detect, and track important biological processes within living cells including protein synthesis, protein glycosylation and nucleic acid proliferation. A limitation of current chemical probes, which have largely focused on mammalian cells, is that they often cannot be applied to other organisms due to metabolic differences. For example, the thymidine derivative 5-ethynyl-2’-deoxyuridine (EdU) is a gold standard metabolic chemical probe for assessing DNA proliferation in mammalian cells however is unsuitable for the study of malaria parasites due to *Plasmodium* species lacking the thymidine kinase enzyme that is essential for metabolism of EdU. Herein we report the design and synthesis of new thymidine-based probes that sidestep the requirement for a thymidine kinase enzyme in *Plasmodium*. Two of these DNADetect™ probes exhibit robust labelling of replicating asexual intraerythrocytic *P. falciparum* parasites, as determined by flow cytometry using copper catalysed azide-alkyne cycloaddition (CuAAC) to a fluorescent azide. The DNADetect™ chemical probes are synthetically accessible and thus can be made widely available to researchers as tools to further understand the biology of different *Plasmodium* species, including laboratory lines and clinical isolates.

## Introduction

A gold standard chemical probe for detection of DNA synthesis and proliferation in mammalian cells is the alkyne modified thymidine analogue 5-ethynyl-2’-deoxyuridine (EdU), which is detected by copper catalysed azide-alkyne cycloaddition (CuAAC) with a fluorescent azide, Figure 1.^1–3^ This reagent exploits the mammalian cell thymidine salvage pathway to incorporate into nuclear DNA. To be effective, EdU requires cellular uptake via nucleoside transporters followed by stepwise phosphorylation with thymidine kinase (TK), thymidylate kinase and nucleoside diphosphate kinase to form a nucleotide triphosphate (designated EdU-P3) followed by integration of the triphosphate into DNA with a DNA polymerase and release of pyrophosphate (PPi),^4^ Figure 1. Detection protocols for EdU in mammalian cells are robust, and commercial reagent kits based on this probe are in widespread use worldwide.^5^

**Figure 1.**
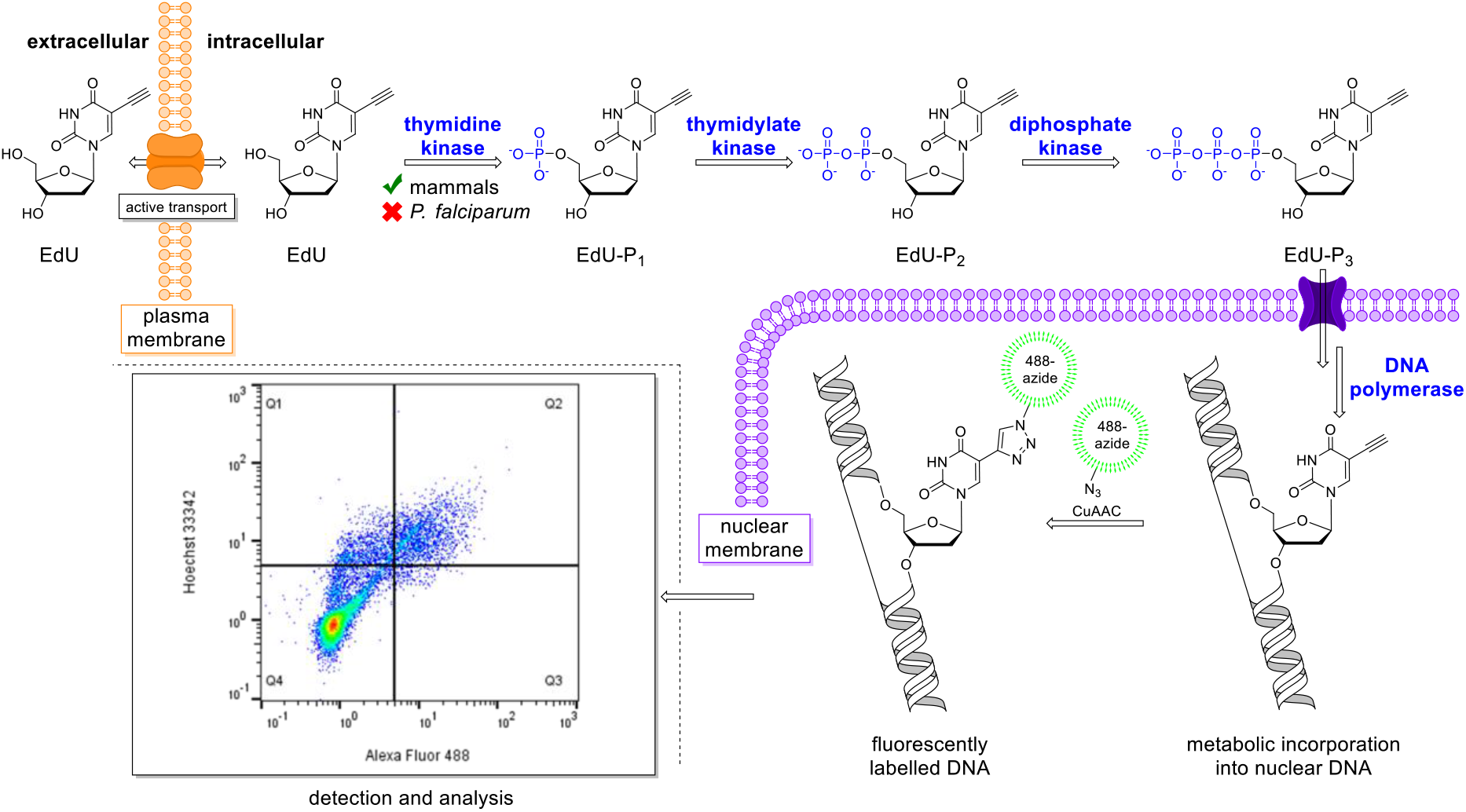
Schematic showing cellular uptake, metabolic processing, and detection of the thymidine derivative EdU when used for detection of S-phase DNA synthesis and proliferation in mammalian cells. *P. falciparum* lacks thymidine kinase, an enzyme that is essential for metabolism of EdU, rendering EdU ineffective for the study of wild type *P. falciparum*.

While used predominantly in mammalian cells, EdU has been applied to study DNA synthesis in pathogens where thymidine metabolism mirrors that occurring in mammalian cells, including intracellular pathogens such as *Trypansoma cruzi* (the causative agent of Chagas disease), *Cryptosporidium parvum* (which causes cryptosporidiosis), and *Naegleria fowleri* (which can cause primary amoebic meningoencephalitis), as well as several diverse unicellular organisms.^6–15^ However, EdU cannot be directly used to detect DNA synthesis in pathogens where differences in thymidine metabolism prevent this, including in *Plasmodium* species which cause the globally important disease malaria. *Plasmodium falciparum* is the most significant of the human infecting *Plasmodium* species and caused an estimated 247 million clinical cases and 619,000 deaths in 2021.^16^ *Plasmodium* lack the thymidine kinase enzyme that phosphorylates thymidine and that is essential for the metabolism of EdU, Figure 1. However, *Plasmodium* parasites do express downstream enzymes in the thymidine biosynthetic pathway needed for eventual incorporation of thymidine into DNA, and both the parasitized host erythrocyte and parasite possess membrane channels that allow nucleoside entry and thus also allow chemical probe uptake.^17–20^ In addition, genetically engineered *P. falciparum* with an introduced promiscuous viral thymidine kinase from *Herpes simplex* virus has been used in DNA labelling studies using BrdU (a thymidine probe developed in the 1970s that requires an antibody for detection) and recently EdU and CuAAC.^21, 22^ Notably, EdU detection requires less harsh processing than BrdU and is a workhorse chemical probe for assessing DNA synthesis during S-phase in mammalian cells and in pathogens that express a thymidine kinase naturally.^23^ While transfection of *P. falciparum* parasites with an exogenous thymidine kinase supports the feasibility of DNA labelling studies using both BrdU and EdU, this approach has limitations, including limiting the flexibility to work with any *Plasmodium* species, such as laboratory lines and field isolates, possible impacts of transgenesis on wildtype *Plasmodium* biology, and biosafety and regulatory compliance considerations associated with using a genetically modified organism. Another chemical probe tool, the alkyne nucleoside purine derivative, 7-deaza-7-ethynyl-2’-deoxyadenosine (EdA), has been shown to label newly synthesized DNA in mammalian cells^24^ and the apicomplexan parasites *Toxoplasma gondii*,^25^ *C. parvum*^25^ as well as *P. falciparum* and *P. vivax* liver stages.^26^ However, while this probe overcomes the need for transfection with an exogenous thymidine kinase in *Plasmodium*, a limitation is that unlike thymidine analogues, purine nucleosides are involved not only in DNA synthesis but in a myriad of other metabolic pathways including cellular energy and intracellular signaling^27^ which may contribute background signals that complicate data interpretation. Furthermore, compared to other proliferation markers, thymidine kinase levels during S-phase have been shown to more accurately determine the rate of DNA synthesis in actively dividing tumour cells.^28^

In this study we have designed next generation metabolic chemical probes for *Plasmodium*, designated DNADetect™, that are designed to bypass the need for thymidine kinase and therefore allow the study of DNA synthesis in wild type parasites. The DNADetect™ probes build on the pedigree of the gold standard EdU probe for labelling of DNA proliferation in mammalian cells while addressing the metabolic hurdles that make EdU ineffective for application in wild type *P. falciparum*.

## Results and Discussion

A panel of compounds that expand on the core structure of EdU was designed to bypass the requirement for thymidine kinase for metabolic incorporation. Specifically, EdU was modified with the addition of a protected phosphate group to the 5’-hydroxyl group of the nucleoside sugar moiety. This design echoes the widely adopted prodrug approach of medicinal chemistry, in particular the use of nucleotide monophosphate prodrugs, known as ‘pronucleotides’.^29^ Pronucleotides are membrane permeable and unmask following cell entry to release the corresponding active nucleoside monophosphate that is trapped and retained intracellularly due to the negatively charged phosphate. Furthermore, the released phosphorylated nucleotide joins the biosynthetic pathway downstream of the first nucleoside kinase, identified as the rate limiting enzyme in the biosynthetic pathway of triphosphates.^29^

Four acyloxybenzyl protected derivatives (compounds **1a-1d**), were designed and synthesized; the new chemical probes are designated DNADetect™, Figure 2a. The variable acyl ester moieties of **1a-1d** were selected to cover a range of lipophilicities and half-lives for release of free EdU monophosphate (EdU-P_1_) to allow comparison of labelling efficacy and includes acetyl (**1a**), benzoyl (**1b**), pivaloyl (**1c**), and heptanoyl (**1d**) acyl esters. Once inside the cell the acyl ester functions as a trigger group that is hydrolyzed, either spontaneously or via nonspecific esterases.^30^ The synthetic installation of the acyloxybenzyl protecting group on EdU was performed by expanding upon phosphoramidite methodology previously optimised by our group (Supporting Information).^8^ In brief, benzyl alcohols **2a-2d** were used to prepare the phosphoramidite reagents **3a-3d**, Figure 2b, which were then used for phosphitylation of EdU to generate the target compounds **1a-1d**, Figure 2c.^30^

**Figure 2.**
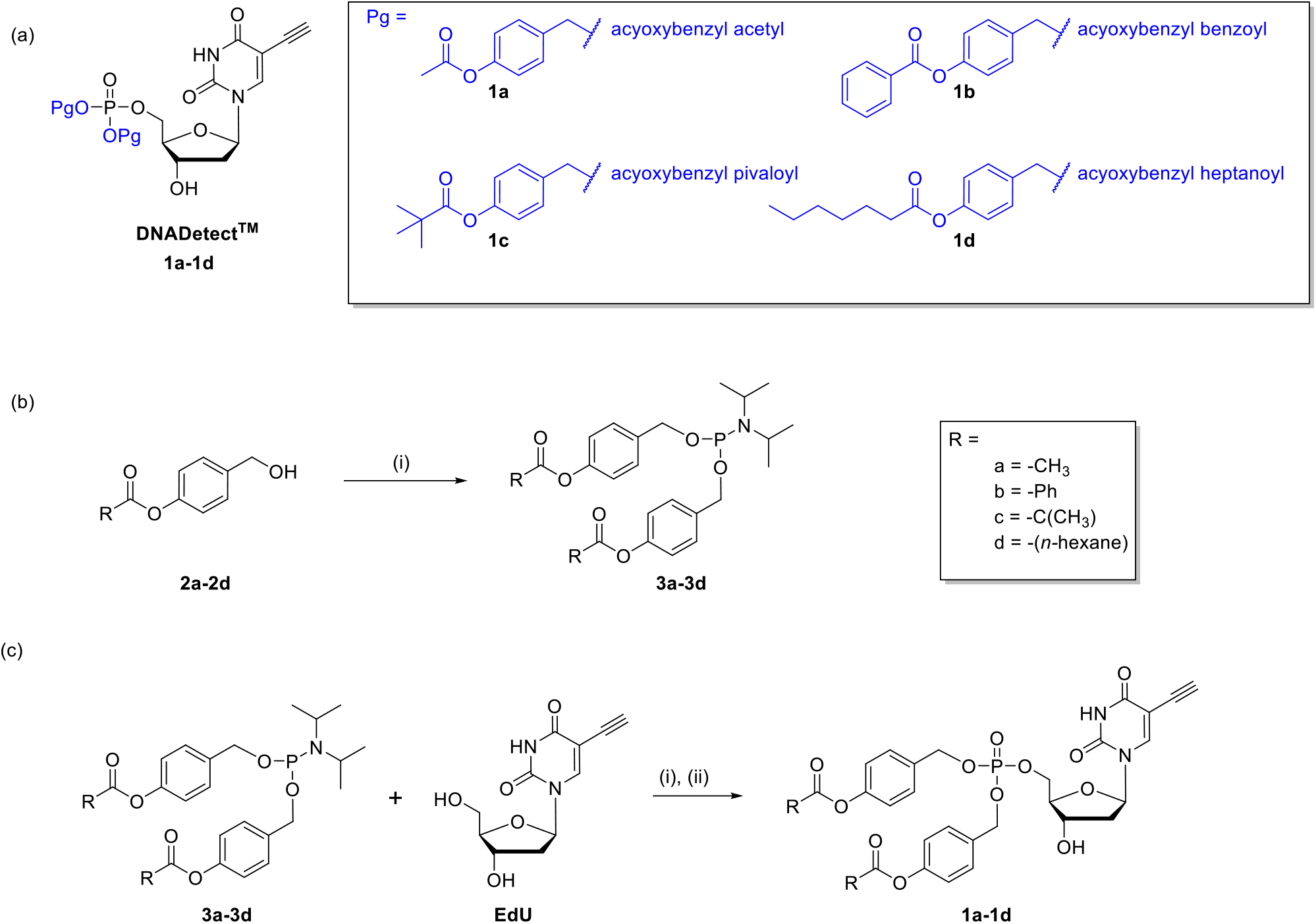
(a) Target chemical probes **1a-1d**. The compounds comprise the core EdU with acyloxybenzyl 5’-phosphate protecting groups (Pg). (b) Synthesis of phosphoramidites **3a-3d** from 4-acyloxybenzylalcohols **2a-2d**: i) PCl_3_ (1 equiv), iPr_2_NH (2 equiv), Et_3_N (2 equiv), THF (Solvent). (c) Phosphitylation of EdU with phosphoramidites **3a-3d** to yield the protected EdU monophosphate chemical probes **1a-1d**: (i) TFA (4 equiv), phosphoramidite (**3a-3d**) pyridine (solvent), CH_2_Cl_2_ (solvent), −20 °C, 30 min. (ii) (1.1 equiv), *t*-BuOOH (5-6 M in decane, 2.4 equiv), pyridine (solvent), CH_2_Cl_2_ (solvent), −5 °C – rt, 30 min.

The mechanism of action of **1a-1d** depends on the interplay of prodrug chemistry, thymidine salvage pathways and enzymatic incorporation into DNA.^31^ The selection of the phosphate protecting group for our intended application is critical as the compounds must have sufficient stability and lipophilicity for parasitized erythrocyte uptake, crossing the erythrocyte membrane, parasitophorous vacuolar membrane and the parasite plasma membrane,^17–20^ and once inside the parasite unmask efficiently to give the free phosphate, EdU-P1, for eventual incorporation of the probe into parasite DNA, Figure 3. Notably, the acyloxybenzyl protecting group has been successfully implemented as a prodrug strategy in *Plasmodium* where it was used to mask the phosphonate group of the antimalarial drug fosmidomycin.^32^ This approved drug provides precedent that *Plasmodium* has capacity to hydrolyse the acyloxybenzyl group.

**Figure 3.**
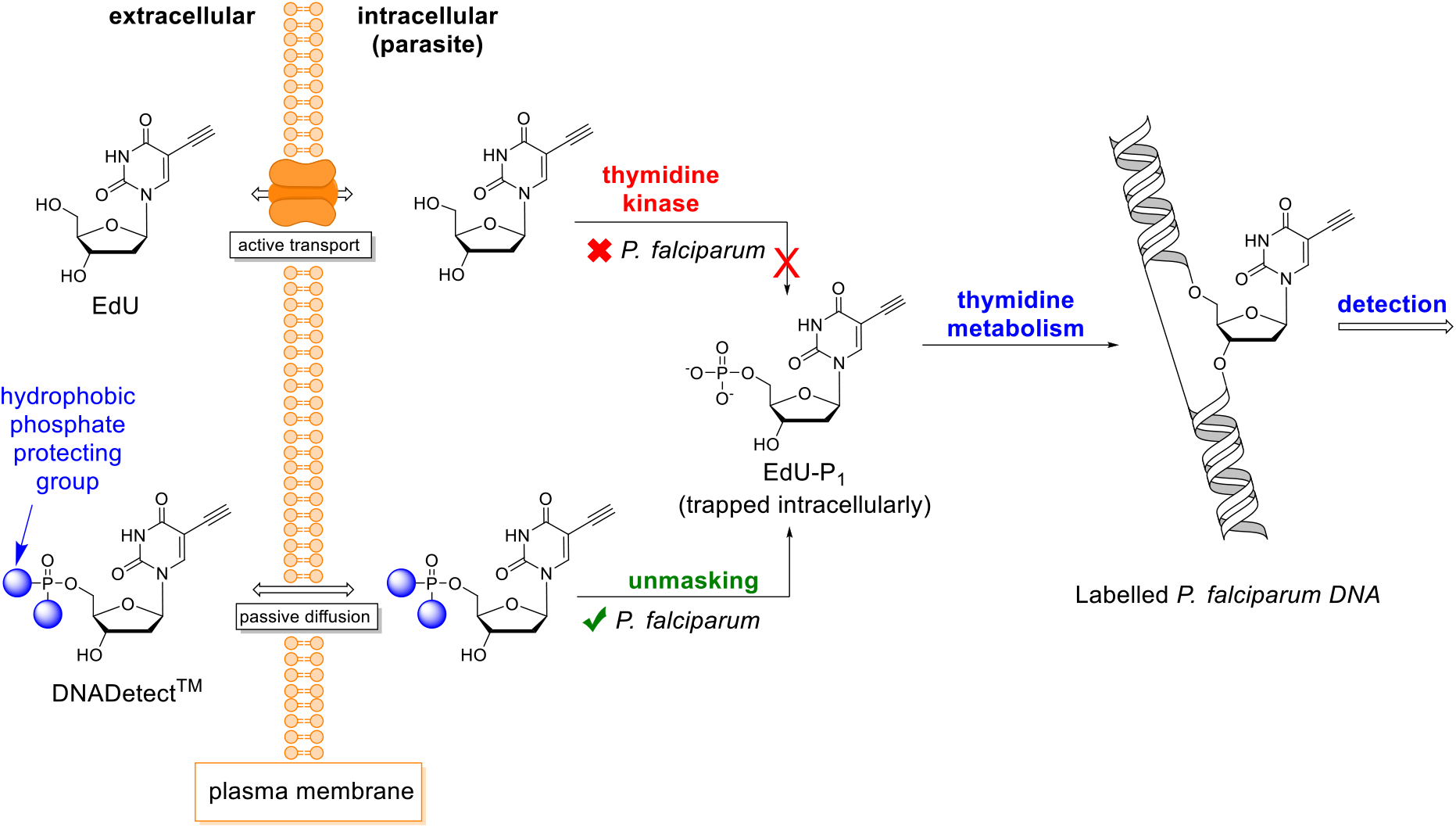
Mechanism of action of DNADetect™ chemical probes designed to bypass the need for thymidine kinase and to allow the detection and study of DNA synthesis in wild type *P. falciparum*.

### *In vitro* growth inhibition activity of probes 1a-1d against *P. falciparum*

To inform selection of concentration windows for labelling DNA of replicating *Plasmodium*, chemical probes **1a-1d** were first assessed for *in vitro* growth inhibition in 72 h dose-response assays against *P. falciparum* 3D7. EdU, EdA and the antimalarial drug chloroquine, to which *P. falciparum* 3D7 is sensitive, were included as controls. As the benzyl alcohol prodrug masking groups **2a-2d** are released as part of the mechanism of action of **1a-1d** that leads to release of the free EdU monophosphate, the benzyl alcohol protecting group controls **2a-2d** were also assessed to ensure they do not display cytotoxicity. The *P. falciparum* 3D7 50% growth inhibition values (IC_50_) of EdU, EdA, protecting group controls (**2a-d**) and compound **1a**, with an acyloxybenzyl acetyl protecting group, demonstrate that these compounds were inactive against *P. falciparum* with IC_50_ > 40 μM, Table 1. Compounds **1c** and **1d**, with acyloxybenzyl pivaloyl and heptanoyl protecting groups, respectively, had low activity against *P. falciparum* (IC_50_ ~10-12 μM; Table 1), while **1b** with an acyloxybenzyl benzoyl protecting group was the most toxic to the parasite under these assay conditions (IC_50_ 1.7 μM; Table 1). As expected,^33, 34^ the antimalarial control drug chloroquine had potent activity (IC_50_ 0.01 μM; Table 1).

**Table 1.**
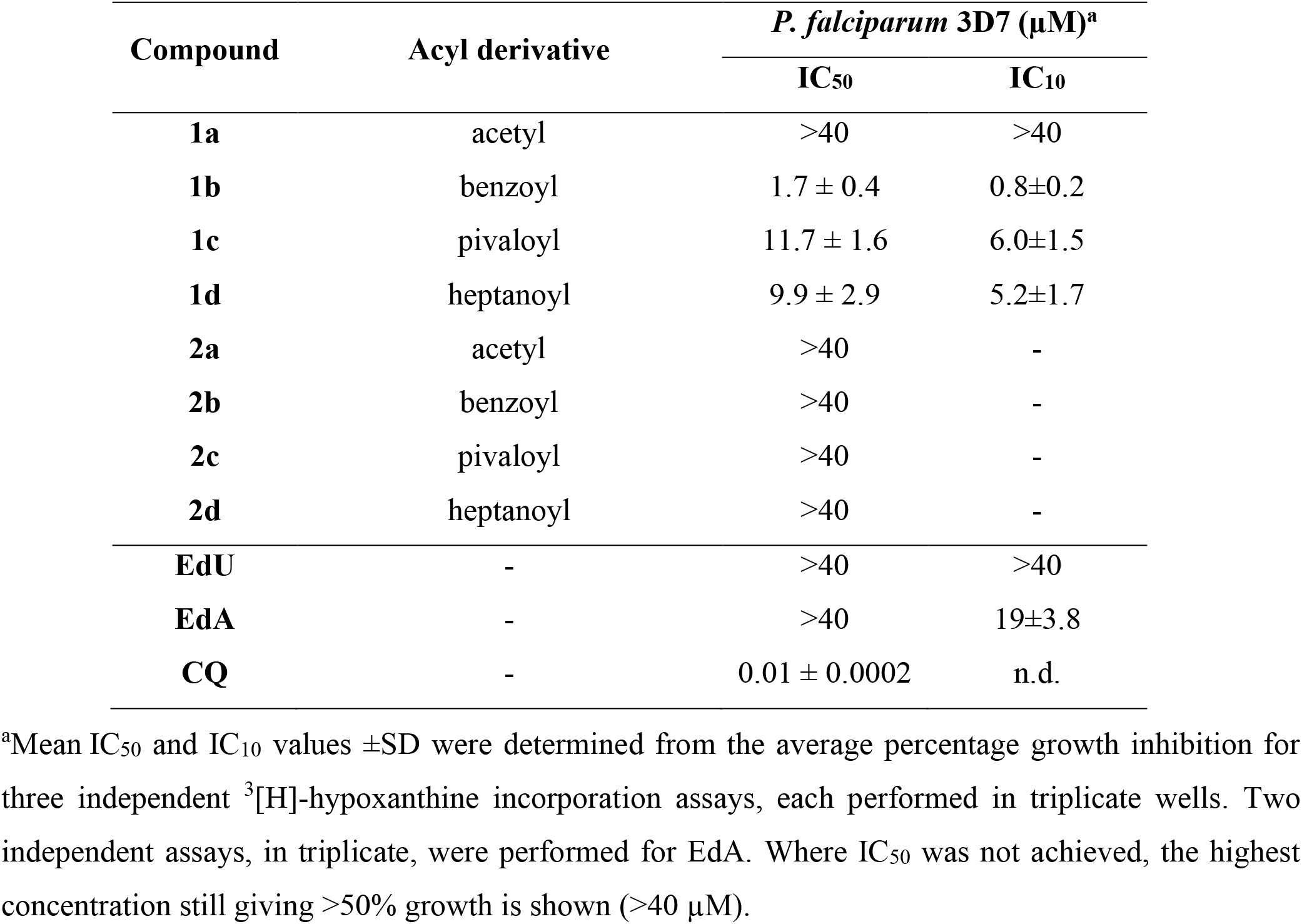
Activity of DNADetect™ probes and controls against *P. falciparum* 3D7.

### Quantitative analysis of *P. falciparum* DNADetect™ probes using flow cytometry

To quantify incorporation of EdU, EdA and **1a-1d** into *P. falciparum* parasitized erythrocytes, flow cytometry was used, Figure 4. Late trophozoite/early schizont stage parasites (~30-40 h post-invasion of merozoites into erythrocytes) were exposed to compound concentrations that inhibited less than 10% of parasite growth (< IC_10_) in 72 h *in vitro* growth inhibition assays (**1b** 0.5 μM; **1c**, 5 μM; **1d**, 5 μM), or for probes with IC_50_ >40 μM at 5 μM (**1a** and EdU). Probe EdA was used at its previously published concentration of 10 μM.^26^ Treatments were performed for 4 h, followed by fixing with 2% paraformaldehyde (Sigma, USA) and 0.2% glutaraldehyde, permeabilization in 0.1% Triton^®^X-100 (Sigma, USA) and a following CuAAC reaction with Alexa Fluor 488 azide to enable flow cytometry detection at 488 nm. Under these conditions, no morphological changes were apparent for any of the treatments, as determined by microscopic examination of QuickDip (POCD, Australia) stained thin blood smears (data not shown). Cells were also treated with Hoechst 33342 nucleic acid stain to differentiate parasitized erythrocytes from uninfected erythrocytes, which are anuclear. The percentage of parasitized erythrocytes that were Alexa Fluor 488 and Hoechst 33342 positive was determined. Approximately 60% of parasitized erythrocytes were Alexa Fluor 488 positive for probe **1d**, with 30% and 20% being Alexa Fluor 488 positive for probes **1c** and **1b**, respectively, Figure 4. In contrast, only ~6% of cells were Alexa Fluor 488 positive for probe **1a**. This level of signal is similar to the negative control EdU^22^ (~3%, Figure 4) and thus is likely to be a non-specific background effect. In contrast to the previous study that reported labeling *P. falciparum* with EdA,^26^ here we observed only background levels (~4%, Figure 4) of Alexa Fluor 488 labelling using this probe. However, as EdA is a purine nucleoside derivative it is possible it is outcompeted by the purines present in the human sera used in the culture media of this study. Given that scavenging of purines from the culture media (e.g. hypoxanthine) is essential for *Plasmodium* parasite nucleic acid replication and growth, this is a known limitation of EdA as a probe for *Plasmodium* DNA replication studies, as previously reported.^26^ These findings indicate that under the labelling conditions used in this study, the heptanoyl acyloxybenzyl protecting group of **1d** is the most efficient in providing a source of EdU-P1 within the parasite to feed the thymidine metabolism pathway for DNA synthesis, followed by the pivaloyl derivative **1c** and benzoyl derivative **1b**.

**Figure 4.**
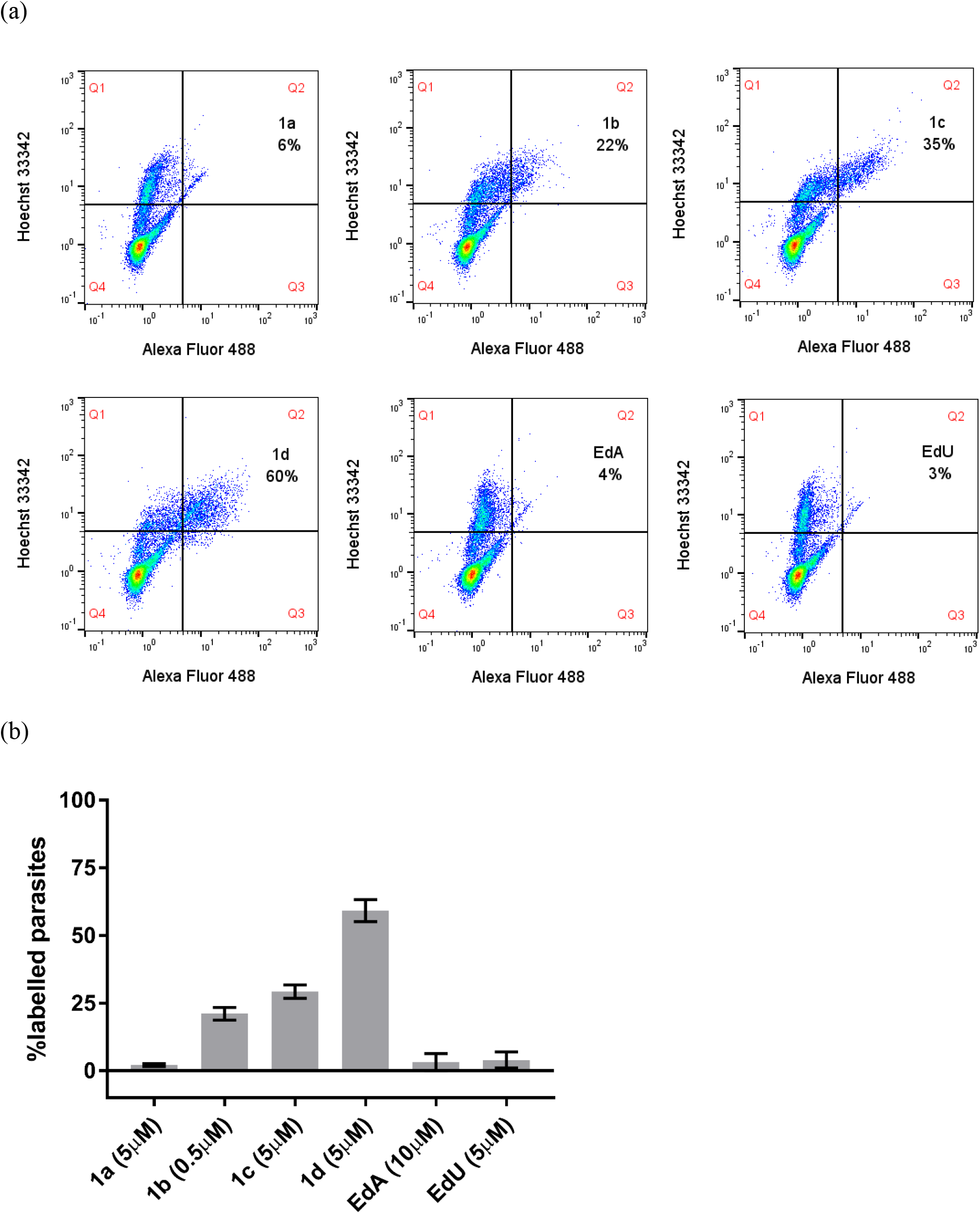
Flow cytometry analysis of incorporation of DNADetect™ probes and controls into *P. falciparum* 3D7. (a) Representative flow cytometry dot plots from one of two independent experiments showing Hoechst 33342 and Alexa Fluor 488 staining of *P. falciparum* 3D7 parasites following 4 h exposure to DNADetect™ probes **1a-1d** and controls EdA and EdU. Population gates are denoted as: Q1, Hoechst 33342 only; Q2, Hoechst 33342 + Alexa Fluor 488; Q3, Alexa Fluor 488 only and Q4, unstained. Percentages annotated in gate Q2 indicate Hoechst 33342 and Alexa Fluor 488 positive parasites. (b) Graph showing labelling efficiency of each probe normalized to the DMSO control presented as mean (± SD) percentage of Alexa Fluor 488 and Hoechst 33342 labelled parasites from at least two independent experiments (20,000 events counted per experiment).

### Analysis of Chemical Probe Stability and Lipophilicity

To be effective in labelling DNA, sufficient intact probe must be available in assay media for cellular uptake. Thus, the stability of **1a-1d** was assessed in *P. falciparum* culture media (RPMI-complete media (Sigma-Aldrich, RPMI 1640 with sodium bicarbonate, 25 mM HEPES, 50 μg/mL hypoxanthine, 5% human serum (Sigma-Aldrich) and 2.5 mg/mL Albumax II (Invitrogen),) and degradation of compounds monitored by LC-MS/MS, Table 2. As expected, increased steric hindrance around the acyl carbonyl of the acyloxybenzyl protecting groups increased the compounds half-life in media with the trend **1c** (pivaloyl; t_1/2_ 1.88 h) >**1b** (benzoyl; t_1/2_ 1.60 h) > **1d** (heptanoyl; t_1/2_ 0.53 h) > **1a** (acetyl; t_1/2_ 0.085 h), yielding stability profiles over a range of an order of magnitude with the pivaloyl derivative **1c** ~ 21-fold more stable than acetyl derivative **1a**, Table 2. These data indicate that the half-life of **1a** is likely too short to be effective as a chemical probe, consistent with flow cytometry data and our observation of low toxicity of probe against *P. falciparum* (>40 μM, Table 1). Compounds **1b-1d** have half-life values sufficiently long to enable uptake into the erythrocytes. The stability trends are similar to those reported for phosphonates bearing the same acyloxybenzyl protecting groups^32^, indicating that the relative stability of these groups in media is a function of the acyl group differences, independent of the underlying common parent compound. To estimate compound lipophilicity, a parameter associated with cell membrane permeability, the cLog P of **1a-1d** was calculated using the Chemaxon chem-bioinformatics software package, Table 2. The cLog P for **1a** was the lowest of the panel (cLog P 2.11), indicative of low cell uptake. The combination of poor uptake with rapid degradation in the assay media is consistent with this compound exhibiting the poorest labelling efficiency of *P. falciparum* parasites (~6%, Figure 4). The cLog P for the pivaloyl derivative **1c** (cLog P 3.94) and benzoyl derivative **1b** (cLog P 4.24) indicate they have lipophilicity that is in the range for good passive membrane permeability for erythrocyte uptake. Compound **1d** with a cLog P of 5.40 is also expected to have good erythrocyte uptake. Compounds **1b-1d** thus have good stability in media and suitable lipophilicity for passive erythrocyte uptake. We therefore suggest that quantitative differences observed in probe **1a-1d** incorporation in flow cytometry are likely due to differences in efficiency of protecting group hydrolysis once inside the malaria parasite.

**Table 2.** Media stability and calculated lipophilicity of DNADetect™ probes **1a-1d**.

| Compound | Half-life in media (h)<sup>a</sup> | cLog P<sup>b</sup> |
| --- | --- | --- |
| **1a** | 0.085 ± 0.004 | 2.11 |
| **1b** | 1.6 ± 0.05 | 4.24 |
| **1c** | 1.88 ± 0.14 | 3.94 |
| **1d** | 0.53 ± 0.03 | 5.40 |

<sup>a</sup>Analysis of 5 µM of compound in RPMI-complete media (Sigma Aldrich, RPMI 1640 with sodium bicarbonate, 25 mM HEPES, 50 µg/mL hypoxanthine, 5% human serum and 2.5 mg/mL Albumax II Invitrogen), incubated at 37 °C in a total volume of 3 mL. <sup>b</sup>Calculated using Chemaxon package for Windows 10.

## Conclusions

The premise of using nucleoside analogues as chemical probes to study DNA synthesis is that these reagents carry a chemical tag that makes newly synthesised DNA distinguishable from the nonreplicating DNA. EdU is the mainstay chemical probe for DNA synthesis in mammalian cells but may not always be directly applied to pathogen studies due to differences in DNA metabolism, including for *Plasmodium* parasites which do not express the thymidine kinase enzyme that is essential for metabolism of EdU. While transgenic *P. falciparum* with an introduced promiscuous thymidine kinase from *Herpes simplex* virus has been successfully used to permit labelling of the parasite with EdU,^22^ this approach is not feasible for many research needs, including the study of non-transgenic *Plasmodium* from different genetic backgrounds, with different natural or selected drug resistance profiles, pathogens from different life cycle stages, different pathogen species (e.g., field isolates which may be difficult to culture *in vitro*) and many other important applications such as dormancy.

Herein we have developed novel chemical probe reagents, DNADetect™, designed to (i) passively cross membranes to enter the *Plasmodium* parasite and (ii) bypass the requirement for thymidine kinase, delivering the downstream EdU monophosphate metabolite that can metabolically incorporate into DNA. We have demonstrated that the DNADetect™ probes **1b-1d** robustly label DNA in *in vitro* cultured wild type asexual stage *P. falciparum* using flow cytometry, and where the current gold standard thymidine probe, EdU, fails due to the absence of thymidine kinase. The new probes provide a complementary solution to expression of exogenous thymidine kinase in the parasite, overcoming limitations associated with transgenesis. Furthermore, as there are other globally significant pathogens that lack thymidine kinase, including *T. gondii* (infects ~30% of humans globally, causes severe morbidity)^35^ and *Mycobacterium tuberculosis* (one of the world’s most lethal infectious diseases.),^36^ we anticipate these novel DNADetect™ probes will enable researchers to study these pathogens where there is a clear benefit and an unmet need for suitable chemical probes for DNA synthesis. We expect that this method will allow high-resolution microscopic or flow cytometry analysis of cellular processes involving DNA, such as dormancy.

## Methods

### *In vitro P. falciparum* growth inhibition assays

*P. falciparum* 3D7^37^ infected erythrocytes were cultured in O positive human erythrocytes in RPMI 1640 media (Gibco, USA) containing 10% heat-inactivated pooled human sera and 5 μg/mL gentamicin (Sigma, USA). Cultures were maintained at 37°C in a gas mixture composed of 5% O2, 5% CO2, and 90% N_2_, as described previously.^38^ *In vitro* inhibition of *P. falciparum* growth inhibition was assessed using a 72 h isotopic microtest, essentially as previously described.^33^ Briefly, highly synchronous ring-stage *P. falciparum* infected erythrocytes obtained by sorbitol treatment^39^ were seeded at 0.25% parasitemia and 2.5% final haematocrit into 96-well tissue culture plates (3596 Costar^®^, Corning, USA) containing serial dilutions of control or test compounds. Compound vehicle only (0.5% final DMSO; also constant in all wells) and the antimalarial drug chloroquine served as negative and positive controls, respectively, in each assay. After incubating for 48 h under standard *P. falciparum* culture conditions, 0.5 μCi [^3^H]-hypoxanthine (PerkinElmer^®^, USA) was added to each well followed by culturing for a further 24 h. Cells were harvested onto 1450 MicroBeta filter mats (Wallac, USA) and ^3^H incorporation determined using a 1450 MicroBeta liquid scintillation counter (PerkinElmer^®^, USA). Percentage inhibition of growth for compound-treated versus matched vehicle only (0.5% DMSO) controls was determined and IC_50_ values calculated using non-linear regression analysis in GraphPad Prism^®^. Each compound was assayed in triplicate wells, in three independent experiments.

### *P. falciparum* flow cytometry assays

*P. falciparum* 3D7 parasites were cultured as described for *in vitro* growth inhibition assays and late trophozoite/early schizont stage infected erythrocytes isolated via Magnetic Activated Cell Sorting (MACS) according to the manufacturer’s instructions using MACS CS columns (Miltenyi, USA). Isolated parasites were resuspended in culture media at 20% parasitaemia and 1% hematocrit and dispensed into 12-well culture plates (1 mL/well) and rested for at least 1 h under standard culture conditions. The cultures were then exposed to test compounds and controls for 4 h at 37 °C at 5 μM (**1a**, **1c**, **1d** and negative control EdU), 0.5 μM (**1b**) or 10 μM (EdA, as previously published^26^). A vehicle only negative control (0.25% DMSO; non-toxic to the parasite) was also included. Following incubation, the cultures were fixed in 2% paraformaldehyde (Sigma, USA) and 0.2% glutaraldehyde (Sigma, USA) for 15 min at room temperature. Cells were then washed twice by resuspending in 1 mL 1% bovine serum albumin (BSA; Bovagen, NZ) in PBS and centrifugation for 1 minute in a mySPIN™6 mini centrifuge (Thermo Scientific, USA). The resulting pellet was permeabilized for 10 min at room temperature in 0.1% Triton^®^X-100 (Sigma, USA) in PBS. After washing twice with 1% BSA/PBS, as above, cell pellets were stained for 30 min at room temperature protected from light, using a CuAAC reaction cocktail (0.2mL) containing 5 μM Alexa Fluor 488 azide (Thermo Fisher, USA), 1 mM CuSO_4_(aq) (Sigma, USA) and 100 mM aqueous sodium ascorbate (Sigma, USA). The cells were then washed twice in 1% BSA/PBS and co-stained with 0.5mL 2 μM Hoechst 33342 (Thermo Fisher, USA) for 15 min at room temperature, protected from light. Following two final washes in 1% BSA/PBS, samples were resuspended in 0.2mL 1% BSA/PBS before acquisition on a MACSQuant^®^ flow cytometer (Miltenyi, USA). Alexa Fluor 488 fluorescence was measured using a blue laser at 488 nm and band pass filter of 525/50 nm. Hoechst 33342 was measured using a violet laser at 405 nm and band pass filter 450/50 nm. The voltage settings and acquisition parameters were as follows, Forward Scatter (FSC) 450v, linear mode; Side Scatter (SSC) 579v, linear mode; Alexa Fluor 488 and Hoechst 33342 400v, hLog. Doublets were excluded by gating on FSC-Area (FSC-A) and FSC-H. Uninfected erythrocytes were included as a background control and 20,000 events were counted per sample. Data were analysed with MACSQuantify software (Miltenyi Biotec, USA) and reported as the mean percentage ±SD (n ≥ 2) of Alexa Fluor 488/Hoechst 33342 positive cells normalized to the DMSO negative control.

### Supporting Information

Additional experimental details, materials, and methods for chemical synthesis and media stability assessment. ^1^H, ^13^C and ^31^P NMR for **1a-1d**. Media stability methods and data analysis.

## Acknowledgments

This research was supported by the Australian Government through the Australian Research Council’s Discovery Projects funding scheme (projects DP220102618, DP180102601). Thank you to the Australian Red Cross Lifeblood for the provision of human blood and sera for culturing *Plasmodium* parasites.

