## Supplementary file for "Thymidine kinase-independent click chemistry DNADetect™ probes for DNA proliferation assessment in malaria parasites"

### Table of Contents

#### Material Page


General Chemistry ........ S3

**Scheme S1:** Synthesis of compounds **3a-d** and **1a-d** ........ S4

General Procedure 1 ........ S5

General Procedure 2 ........ S5

Purification Method A ........ S5

Purification Method B ........ S6

Synthesis of **3a** (Bis(4-acetyloxybenzyl)-*N*,*N*-diisopropylphosphoramidite) ........ S6

Synthesis of **3b** (Bis(4-benzoyloxybenzyl)-*N*,*N*-diisopropylphosphoramidite) ........ S6

Synthesis of **3c** (Bis(4-*tert*-butyloxybenzyl)-*N*,*N*-diisopropylphosphoramidite) ........ S6

Synthesis of **3d** (Bis(4-heptanoyloxybenzyl)-*N*,*N*-diisopropylphosphoramidite) ........ S6

Synthesis of **1a** (Bis(4-acetyloxybenzyl)-EdU monophosphate) ........ S7

Synthesis of **1b** (Bis(4-benzoyloxybenzyl)-EdU monophosphate) ........ S7

Synthesis of **1c** (Bis(4-*tert*-butyloxybenzyl)-EdU monophosphate) ........ S8

Synthesis of **1d** (Bis(4-heptanoyloxybenzyl)-EdU monophosphate) ........ S9

Compound **1a** $^{1}$H and $^{13}$C{H} NMR (DMSO-$d_6$) ........ S10

Compound **1a** $^{31}$P NMR (DMSO-$d_6$) ........ S11

Compound **1b** $^{1}$H and $^{13}$C{H} NMR (DMSO-$d_6$) ........ S12

Compound **1b** $^{31}$P NMR (DMSO-$d_6$) ........ S13

Compound **1c** $^{1}$H and $^{13}$C{H} NMR (DMSO-$d_6$) ........ S14

Compound **1c** $^{31}$P{H} NMR (DMSO-$d_6$) ........ S15

Compound **1d** $^{1}$H and $^{13}$C{H} NMR (DMSO-$d_6$) ........ S16

Compound **1d** $^{31}$P{H} NMR (DMSO-$d_6$) ........ S17

Media stability studies of chemical probes **1a-1d** ........ S18

**Table S1:** Mass transitions summed for compounds **1a-1d** ........ S20

**Table S2:** Degradation of compounds **1a-1d** in RPMI-complete media at 37 °C ........ S20

**Figure S1:** Stability of compounds **1a-1d** in RPMI-complete media at 37 °C ........ S21

**References** ........ S22

#### General Chemistry

All reactions were carried out in dry solvents under anhydrous conditions unless otherwise stated. All chemicals were purchased from commercial suppliers and used without further purification. All reactions were monitored by TLC using silica plates with visualisation of eluted bands by UV fluorescence ($\lambda$= 254 nm) and charring with vanillin stain (6 g vanillin in 100 mL of EtOH containing 1% v/v 98% Sulfuric acid. Silica gel flash chromatography was performed using silica gel 60 Å (230-400 mesh) or alumina 60 Å (70-230 mesh), where specified. NMR ($^{1}$H, $^{13}$C, $^{19}$F, COSY, NOESY, HSQC and HMBC) spectra were recorded on a Bruker AVANCE III HD 500 MHz NMR spectrometer equipped with a BBO probe at 25 °C. Chemical Shifts for $^{1}$H and $^{13}$C NMR obtained in DMSO-$d_6$ are reported in ppm relative to residual solvent proton (d = 2.50 ppm) and carbon (d = 39.5 ppm) signals, respectively. Chemical Shifts for $^{1}$H and $^{13}$C NMR obtained in $CDCl_3$ are reported in ppm relative to residual solvent proton (d = 7.26 ppm) and carbon (d = 77.16 ppm) signals, respectively. The $^{31}$P NMR chemical shifts are reported in ppm using neat $(EtO)_3P$ (-1 ppm) as an external reference, employing coaxial insert tubes. Signal splitting multiplicity is indicated as follows: s (singlet), d (doublet), t (triplet), q (quartet), m (multiplet), dd (doublet of doublets), br (broad signal), a (apparent). Assignments of $^{1}$H and $^{13}$C chemical shifts were established by COSY, HSQC, HMBC, and NOESY experiments. Coupling constants are reported in hertz (Hz). Low resolution mass spectrometry (LRMS) data were acquired on a Thermo Fisher MSQ Plus single quadruple ESI mass spectrometers using electrospray as the ionisation technique in positive and/or negative mode as stated. High resolution mass spectrometry (HRMS) data were acquired on a 12 T SolariX XR FT-ICR-MS using electrospray as the ionisation technique in positive-ion and/or negative mode as stated. All MS analysis samples were prepared as solutions in either methanol or acetonitrile. Purity of the compounds were >95% as determined by Thermo Fisher Dionex Ultimate 3000 series HPLC via UV detection at 254 nm. The melting points are uncorrected. 4-acyloxybenzyl alcohols; **2a** (4-acetyloxybenzyl alcohol), **2b** (4-benzoyloxybenzyl alcohol), **2c** (4-*tert*-butyloxybenzyl alcohol), **2d** (4-heptanoyloxybenzyl alcohol) were synthesised according to literature procedures.[9, 10]

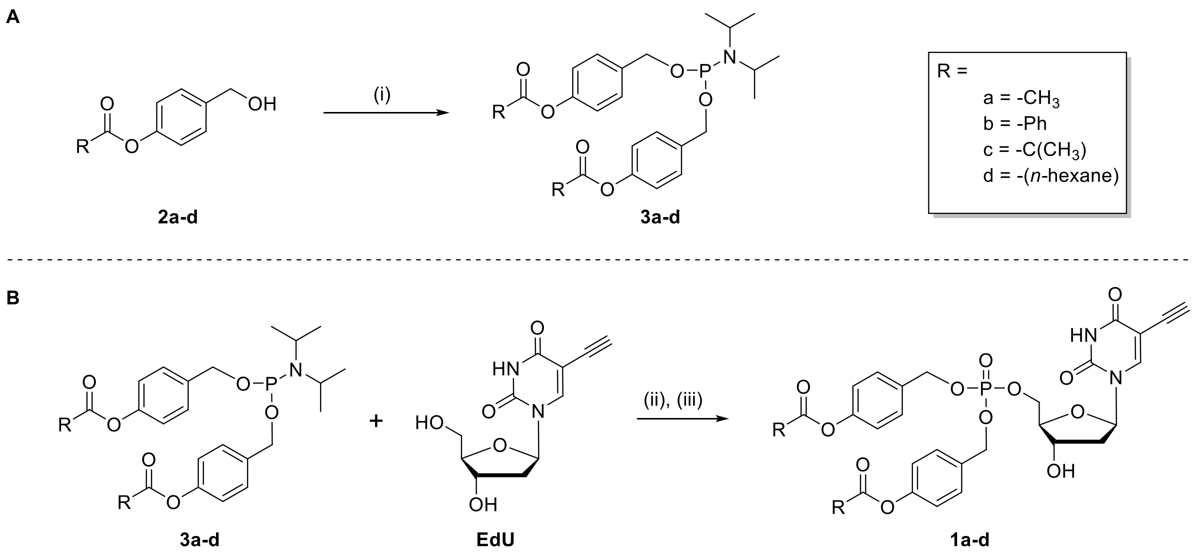


**Scheme S1.** Synthesis of compounds **3a-d** and **1a-**d. A) synthesis of phosphoramidites **3a-d** from 4-acyloxybenzylalcohols. i) $PCl_3$ (1 equiv), $iPr_2NH$ (2 equiv), $Et_3N$ (2 equiv), THF (Solvent). **B**) phosphitylation of EdU employing phosphoramidites **3a-d** to yield EdU monophosphate prodrugs. ii) TFA (4 equiv), phosphoramidite (3**a-d**) pyridine (solvent), $CH_2Cl_2$ (solvent), -20 °C, 30 min. (iii) (1.1 equiv), *t*-BuOOH (5-6 M in decane, 2.4 equiv), pyridine (solvent), $CH_2Cl_2$ (solvent), -5 °C – rt, 30 min.

**General procedure 1**. **Phosphitylation of free nucleoside analogues with phosphoramidites.**

A solution of the nucleoside derivative in pyridine was dried over 3Å molecular sieves (3Å - MS) for a minimum of 16 h at rt. TFA (4 equiv) was added and the mixture left to stand for at least one hour. The mixture was then removed from the 3Å-MS by syringe, added to a round bottom-flask and cooled to -20 °C. The phosphoramidite (1.1 equiv) was dissolved in a minimum amount of $CH_2Cl_2$ (4 equiv) and added to the nucleoside substrate slowly with vigorous stirring. The mixture was allowed to warm to RT over 30 min, at which point the mixture was cooled to -5 °C and *t*-BuOOH (5-6 M in decane, 2.4 equiv) was added and the mixture allowed to again warm to rt over 30 min. The reaction mixture was concentrated *in vacuo* and the residue dissolved in MeOH (10 mL). Amberlite IR-120 $H^+$ resin (prewashed MeOH and dried) and Amberlite IRA-21 resin (prewashed with MeOH and dried) were combined in a 1:2 ratio, respectively, then mixed by vortexing. The mixed resins were then added to the MeOH solution of crude product and stirred until pyridine was no longer detectible by TLC (UV) and universal pH indicator paper indicated neutral pH. The mixture was then filtered, and the resin washed with MeOH (10 mL). The filtrate was concentrated *in vacuo* and the crude product purified by silica gel flash chromatography as described for each compound.

**General procedure 2. Synthesis of phosphoramidite reagents.**

$PCl_3$ (1 equiv), was dissolved in $Et_2O$ and cooled to -78 °C then $iPr_2NH$ (2 equiv) and $Et_3N$ (2 equiv) were added to the reaction mixture slowly. The mixture was allowed to warm to 0 °C over the course of 2 h. The mixture was cooled again to -78 °C and the substrate benzyl alcohol **2a-2d** was dissolved in THF then added with vigorous stirring. After stirring for 2 h and warming to 0 °C, the precipitated amine salts were removed by filtration under a $N_2$ atmosphere using a Schlenk filter and with the receiver flask cooled to 0 °C. The resulting cloudy white solution was then concentrated *in vacuo* without heating. The resulting oil was purified employing the methods as stated for each compound.

**Purification Method A**: A short alumina chromatography column (plug) was prepared from basic alumina which had previously been dried at 150 °C for 16 h and then conditioned with $CH_2Cl_2$ (stabilised with 0.1 % EtOH, acid free). The desired fractions of purified compound were pooled together then concentrated *in vacuo* without heating and stored at –20 °C.

**Purification Method B**: The compound was purified by silica gel flash chromatography with 5% triethylamine and 95% *n*-hexanes as mobile phase. The desired fractions were pooled together then concentrated *in vacuo* without heating and stored at –20 °C.

**Bis(4-acetyloxybenzyl)-*N,N*-diisopropylphosphoramidite 3a:** Compound **3a** was prepared from **2a** (4-acetyloxybenzyl alcohol) (0.400 g, 2.407 mmol) according to **General procedure 2 and Purification Method A.** The title compound was afforded as an oil (0.425 g, 85%). The NMR spectroscopic data was consistent with literature values.[1] **$^1$H NMR (500 MHz, $CDCl_3$)** δ 7.35 (d, *J* = 8.3 Hz, 4H), 7.08 – 7.01 (m, 4H), 4.78 – 4.63 (m, 4H), 3.69 (dp, *J* = 10.0, 6.8 Hz, 2H), 1.20 (d, *J* = 6.9 Hz, 12H).

**Bis(4-benzoyloxybenzyl)-*N,N*-diisopropylphosphoramidite 3b**: Compound **3b** was prepared from **2b** (4-benzoyloxybenzyl alcohol**)** (0.400 g, 1.752 mmol) according to **General Procedure 2 and Purification Method A.** The title compound was afforded as an oil (0.410 g, 80%). The NMR spectroscopic data was consistent with literature values.[1] **$^1$H NMR (500 MHz, $CDCl_3$)** δ 8.26 – 8.16 (m, 4H), 7.72 – 7.58 (m, 2H), 7.51 (ddt, *J* = 7.7, 6.5, 1.1 Hz, 4H), 7.46 – 7.38 (m, 4H), 7.22 – 7.14 (m, 4H), 4.85 – 4.68 (m, 4H), 3.72 (dp, *J* = 10.1, 6.8 Hz, 2H), 1.23 (d, *J* = 6.8 Hz, 12H).

**Bis(4-*tert*-butyloxybenzyl)-*N,N*-diisopropylphosphoramidite 3c**: Compound **3c** was prepared from **2c** (4-*tert*-butyloxybenzyl alcohol) (0.500 g, 2.401 mmol) according to **General Procedure 2 and Purification Method A.** The title compound was afforded as an oil (0.444 g, 68%). The NMR spectroscopic data was consistent with literature values.[1] **$^1$H NMR (500 MHz, $CDCl_3$)** δ 7.37 – 7.31 (m, 4H), 7.04 – 6.98 (m, 4H), 4.79 – 4.63 (m, 4H), 3.77 – 3.61 (m, 2H), 1.35 (s, 16H), 1.20 (d, *J* = 6.8 Hz, 14H).

**Bis(4-heptanoyloxybenzyl)-*N,N*-diisopropylphosphoramidite 3d**: Compound **3d** was prepared from **2d** (4-heptanoyloxybenzyl alcohol) (0.500 g, 2.115 mmol) according to **General Procedure 2 and Purification B.** The title compound was afforded as oil (0.424 g, 67%). The NMR spectroscopic data was consistent with literature values.[2] **$^1$H NMR (500 MHz, $CDCl_3$)** δ 7.38 – 7.31 (m, 4H), 7.06 – 7.01 (m, 4H), 4.83 – 4.61 (m, 4H), 3.68 (dp, *J* = 10.1, 6.8 Hz, 2H), 2.54 (t, *J* = 7.5 Hz, 4H), 1.75 (p, *J* = 7.5 Hz, 4H), 1.45 – 1.38 (m, 4H), 1.34 (tdd, *J* = 7.3, 4.6, 3.5 Hz, 8H), 1.20 (d, *J* = 6.8 Hz, 12H), 0.94 – 0.86 (m, 6H).

**Bis(4-acetyloxybenzyl)-EdU monophosphate 1a**:

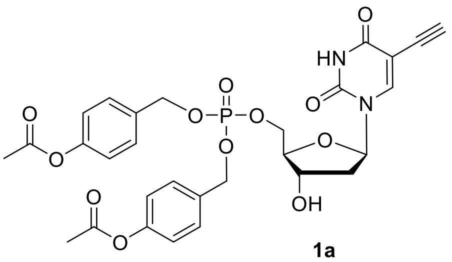

1a

Chemical Formula: $C_{29}H_{29}N_2O_{12}P$

Compound **1a** was synthesised from **EdU** (0.0650 g, 0.2577 mmol) according to **General Procedure 1** employing **3a**. The crude product was purified by silica gel flash chromatography (5% MeOH in $CH_2Cl_2$) to afford the title compound as a colourless gum (0.0410 g, 26%). *Rf* 0.55 (10% MeOH in $CH_2Cl_2$). **$^1$H NMR (500 MHz, DMSO-*d*$_6$)** δ 11.68 (s br, 1H, NH), 7.97 (s, 1H, H-6), 7.44 – 7.39 (m, 4H, 4× H-Ar$_{ortho}$), 7.16 – 7.11 (m, 4H, 4× H-Ar$_{meta}$), 6.11 (t, *J* = 6.9 Hz, 1H, H-1′), 5.43 (d, *J* = 4.4 Hz, 1H, OH-3′), 5.05 (d, *J* = 8.1 Hz, 4H, 2× BnCH$_2$), 4.25 – 4.12 (m, 3H, H-3′, H-5′ and H-5″ ), 4.11 (s, 1H, C≡CH), 3.98 – 3.90 (m, 1H, H-4′), 2.27 (s, 6H, 2× AcO), 2.20 – 2.07 (m, 2H, H-2′$_{(\alpha \text{ and } \beta)}$). **$^{13}$C{H} NMR (125 MHz, DMSO-*d*$_6$)** δ 169.2 (2× *C*O$_2$CH$_3$), 161.7 (C-4), 150.5 (2× C-Bn$_{para}$), 149.5 (C-2), 144.2 (C-6), 133.46 (d, $^3J_{CP}$ = 6.8 Hz, 2× C-Bn$_{ipso}$), 129.2 (4× C-Bn$_{ortho}$), 121.9 (4× C-Bn$_{meta}$), 98.0 (C-5), 85.0 (C-1′), 84.68 (d, $^3J_{CP}$ = 7.4 Hz, C-4′), 83.9 (C≡*C*H), 76.2 (*C*≡CH), 69.9 (C-3′), 68.2 (m, 2× Bn*C*H$_2$), 66.95 (d, $^2J_{CP}$ = 5.4 Hz, C-5′), 39.0 (C-2′ coincident with DMSO solvent peak), 20.9 (2× CO$_2$*C*H$_3$). **$^{31}$P{H} NMR (202 MHz, DMSO-*d*$_6$)** δ -0.75. ***m/z* (LRMS ESI)** 629 [M+H]$^+$, 627 [M-H]$^-$. ***m/z* (HRMS ESI$^+$)** [M + Na]$^+$ found 651.1344; calcd for $C_{29}H_{29}N_2NaO_{12}P$ 651.1350.

**Bis(4-benzoyloxybenzyl)-EdU monophosphate 1b**:

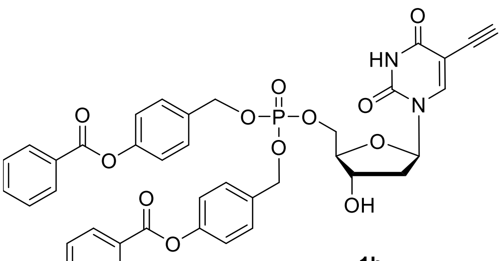

1b

Chemical Formula: $C_{39}H_{33}N_2O_{12}P$

Compound **1b** was synthesised from **EdU** (0.0500 g, 0.1982 mmol) according to **General Procedure 1** employing **3b**. The crude product was purified by silica gel flash chromatography (5% MeOH in $CH_2Cl_2$) to afford the title compound as a colourless gum (0.0504 g, 34%). *Rf* 0.6 (10% MeOH in $CH_2Cl_2$). **$^1$H NMR (500 MHz, DMSO-*d*$_6$)** δ 11.69 (s, 1H, NH), 8.22 – 8.06 (m, 4H, 4× H-Bz$_{ortho}$), 7.98 (s, 1H, H-6), 7.81 – 7.69 (m, 2H, 2× H-Bz$_{para}$), 7.67 – 7.55 (m, 4H, 4× H-Bz$_{meta}$), 7.54 – 7.44 (m, 4H, 4× H-Bn$_{ortho}$), 7.38 – 7.26 (m, 4H, 4× H-Bn$_{meta}$), 6.14 (t, *J* = 6.8 Hz, 1H, H-1′), 5.45 (d, *J* = 4.3 Hz, 1H, OH-3′), 5.11 (d, *J* = 8.2 Hz, 4H, 2× BnCH$_2$), 4.31 – 4.14 (m, 3H, H-3′ and H-5′ and 5″), 4.10 (s, 1H, C≡CH), 3.97 (dd, *J* = 4.2 Hz, 1H, H-4′), 2.25 – 2.09 (m, 2H, H-2′$_{(\alpha \text{ and } \beta)}$). **$^{13}$C{H} NMR (125 MHz, DMSO-*d*$_6$)** δ 164.5

(2× Ar*C*$O_2$-), 161.6 (C-4), 150.6 (2× Ar-O), 149.4 (C-2), 144.2 (C-6), 134.1 (2× C-$Bz_{para}$), 133.7 (d, *J* = 7.5 Hz, 2× C-$Bn_{ipso}$), 129.8 (4× C- $Bz_{meta}$), 129.3 (4× C-$Bn_{ortho}$), 129.0 (4× C-$Bz_{ortho}$), 128.8 (2× C-$Bz_1$), 122.0 (4× C-$Bn_{meta}$), 98.0 (C-5), 85.0 (C-1′), 84.70 (d, *J* = 7.0 Hz, C-4′), 83.8 (C≡*C*H), 76.3 (*C*≡CH), 69.9 (C-3′), 68.22 – 68.10 (m, 2× Bn*C*$H_2$), 67.45 (d, *J* = 5.1 Hz, C-5′), 39.4 (C-2′ coincident with DMSO solvent peak). **$^{31}$P{H} NMR (202 MHz, DMSO-*$d_6$*)** δ -0.68. ***m/z* (LRMS ESI)** 647 $[M+H]^+$ , 645 $[M-H]^-$. ***m/z* (HRMS ESI$^+$)** $[M + Na]^+$ found 775.1656; calcd for $C_{39}H_{33}N_2NaO_{12}P$ 775.1663.

**Bis(4-tert-butyloxybenzyl)-EdU monophosphate 1c**:

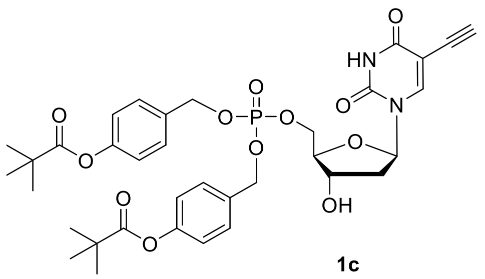

**1c**

Chemical Formula: $C_{35}H_{41}N_2O_{12}P$

Compound **1c** was synthesised from **EdU** (0.0650 g, 0.2577 mmol) according to **General Procedure 1** employing **3c**. The crude product was purified by silica gel flash chromatography (5% MeOH in $CH_2Cl_2$) to afford the title compound as a colourless gum (0.0762 g, 41%). *Rf* 0.37 (5% MeOH in $CH_2Cl_2$). **$^1$H NMR (500 MHz, DMSO-*$d_6$*)** δ 11.68 (s, 1H, NH), 7.96 (s, 1H, H-6), 7.53 – 7.29 (m, 4H, 4× $Bn_{ortho}$), 7.23 – 6.96 (m, 4H, 4× $Bn_{meta}$), 6.12 (t, *J* = 6.8 Hz, 1H, H-1′), 5.50 – 5.38 (m, 1H, OH-3′), 5.05 (d, *J* = 8.2 Hz, 4H, Bn$CH_2$), 4.27 – 4.12 (m, 3H, H-5′, H-5″, H-3′), 4.09 (s, 1H, C≡CH), 3.95 (q, *J* = 4.0 Hz, 1H, H-4′), 2.22 – 2.03 (m, 2H, H-2′$_{(\alpha \text{ and } \beta)}$), 1.30 (s, 18H, 2× $(CH_3)_3CCO_2$). **$^{13}$C{H} NMR (125 MHz, DMSO-*$d_6$*)** δ 176.4 (2× $(CH_3)_3C$*C*$O_2$), 161.6 (C-4), 150.8 (2× $Bn_{para}$), 149.4 (C-2), 144.2 (C-6), 133.44 (dd, *J* = 6.7, 1.3 Hz, 2× $Bn_{ipso}$), 129.20 (d, *J* = 1.7 Hz, 2× $Bn_{ortho}$), 121.8 (2× $Bn_{meta}$), 98.1 (C-5), 85.0 (C-1′), 84.71 (d, *J* = 7.5 Hz, C-4′), 83.9 (C≡*C*H), 76.2 (*C*≡CH), 69.9 (C-3′), 68.19 (d, *J* = 4.7 Hz, 2× Bn$CH_2$), 66.94 (d, *J* = 5.2 Hz, C-5′), 39.0 (C-2′), 38.6 ($(CH_3)_3$*C*$CO_2$) 26.7 ((*C*$H_3)_3CCO_2$). **$^{31}$P{H} NMR (202 MHz, DMSO-*$d_6$*)** δ -0.81. ***m/z* (LRMS ESI)** 713 $[M+H]^+$, 711 $[M-H]^-$. ***m/z* (HRMS ESI$^+$)** $[M + Na]^+$ found 735.2293; calcd for $C_{35}H_{41}N_2NaO_{12}P$ 735.2289.

**Bis(4-heptanoyloxybenzyl)-EdU monophosphate 1d**:

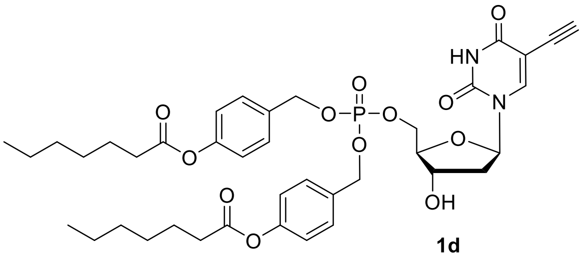

**1d**

Chemical Formula: $C_{39}H_{49}N_2O_{12}P$

Compound **1d** was synthesised from **EdU** (0.0650 g, 0.2577 mmol) according to **General Procedure 1** employing **3d**. The crude product was purified by silica gel flash chromatography (5% MeOH in $CH_2Cl_2$) to afford the title compound as a colourless gum (0.0634g, 32% yield). *Rf* 0.34 (5% MeOH in $CH_2Cl_2$). **$^1$H NMR (500 MHz, DMSO-*d*$_6$)** δ 11.68 (s, 1H, NH), 7.96 (s, 1H, H-6), 7.47 – 7.35 (m, 4H, 4× $Bn_{ortho}$), 7.11 (dq, *J* = 7.7, 2.8 Hz, 4H, 4× $Bn_{meta}$), 6.11 (t, *J* = 6.8 Hz, 1H, H-6′), 5.44 (d, *J* = 4.4 Hz, 1H, OH-3′), 5.04 (d, *J* = 8.2 Hz, 4H, 2× $BnCH_2$ ), 4.25 – 4.11 (m, 3H, H-5′, H-5″, H-3′), 4.09 (s, 1H, C≡CH), 3.94 (dd, *J* = 3.9 Hz, 2H, H-4′), 2.57 (t, *J* = 7.4 Hz, 4H, 2× α-$CH_2$ heptanoyl), 2.22 – 2.06 (m, 2H, H-2′$_{(\alpha \text{ and } \beta)}$), 1.63 (p, *J* = 7.4 Hz, 4H, 2× β-$CH_2$ heptanoyl), 1.42 – 1.25 (m, 12H, 2× γ-$CH_2$, 2× ε-$CH_2$, 2× δ-$CH_2$ heptanoyl), 0.95 – 0.82 (m, 6H, (2× ζ-$CH_3$ heptanoyl). **$^{13}$C{H} NMR (125 MHz, DMSO-*d*$_6$)** δ 171.8 ($CO_2$ heptanoyl), 161.5 (C-4), 150.5 ($Bn_{para}$), 149.4 (C-2), 144.2 (C-6), 133.43 (d, *J* = 6.5 Hz), 129.2 (4× $Bn_{ortho}$), 121.9 (4× $Bn_{meta}$), 98.1 (C-5), 85.0 (H-6′), 84.71 (d, *J* = 7.4 Hz, H-4′), 83.9 (C≡*C*H), 76.1 (*C*≡CH), 69.9 (C-3′), 68.25 – 68.09 (m, 2× Bn*C*$H_2$), 66.94 (d, *J* = 4.6 Hz, C-5′), 39.0 (C-2′), 33.5 (2× α-$CH_2$ heptanoyl), 30.9 (2× δ-$CH_2$ heptanoyl), 28.1 (2× γ-$CH_2$ heptanoyl), 24.3 (2× β-$CH_2$ heptanoyl), 22.0 (2× ε-$CH_2$ heptanoyl), 13.9 (2× ζ-$CH_3$ heptanoyl). **$^{31}$P{H} NMR (202 MHz, DMSO-*d*$_6$)** δ -0.85. ***m/z* (LRMS ESI)** 769 $[M+H]^+$, 767 $[M-H]^-$. ***m/z* (HRMS ESI$^+$)** $[M + Na]^+$ found 791.2915; calcd for $C_{39}H_{49}N_2NaO_{12}P$ 791.2920.

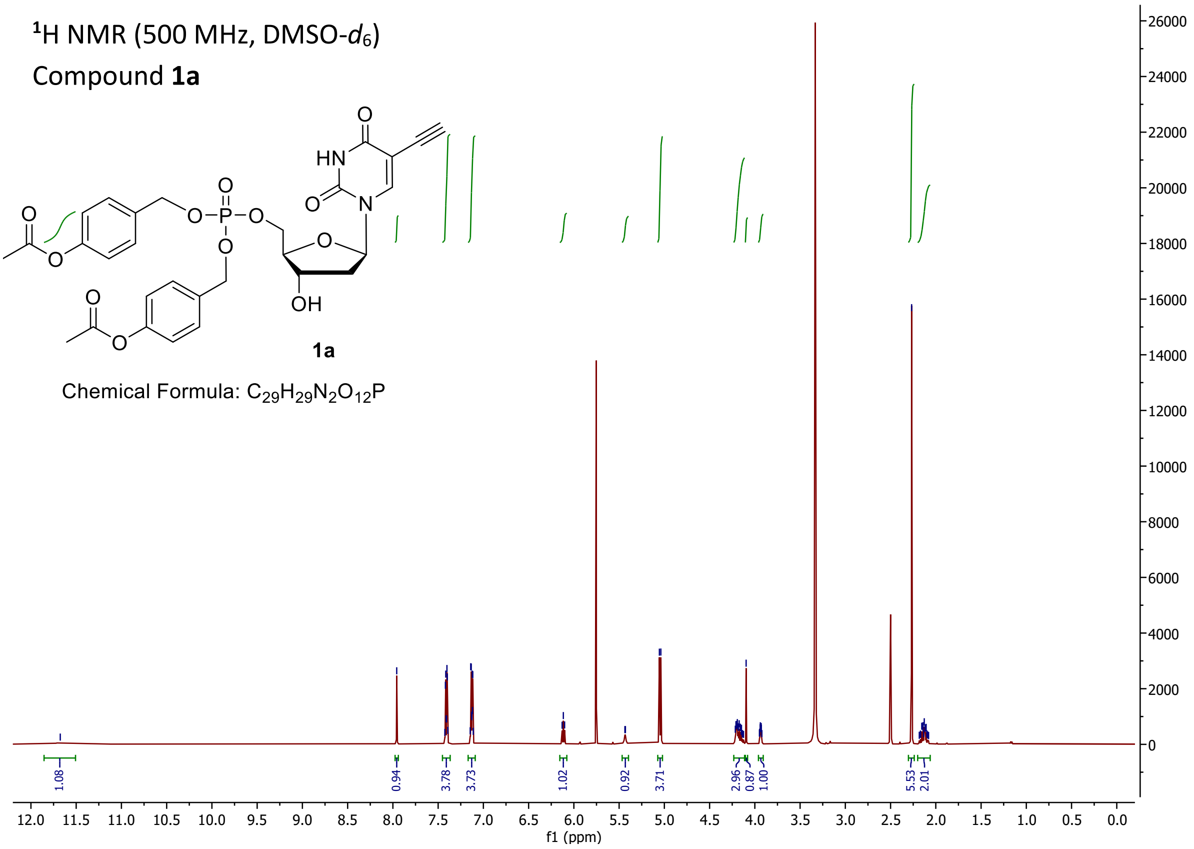

1H NMR (500 MHz, DMSO-d6)
Compound 1a
1a
Chemical Formula: C29H29N2O12P
1.08
0.94
3.78
3.73
1.02
0.92
3.71
2.96
0.87
1.00
5.53
2.01
f1 (ppm)


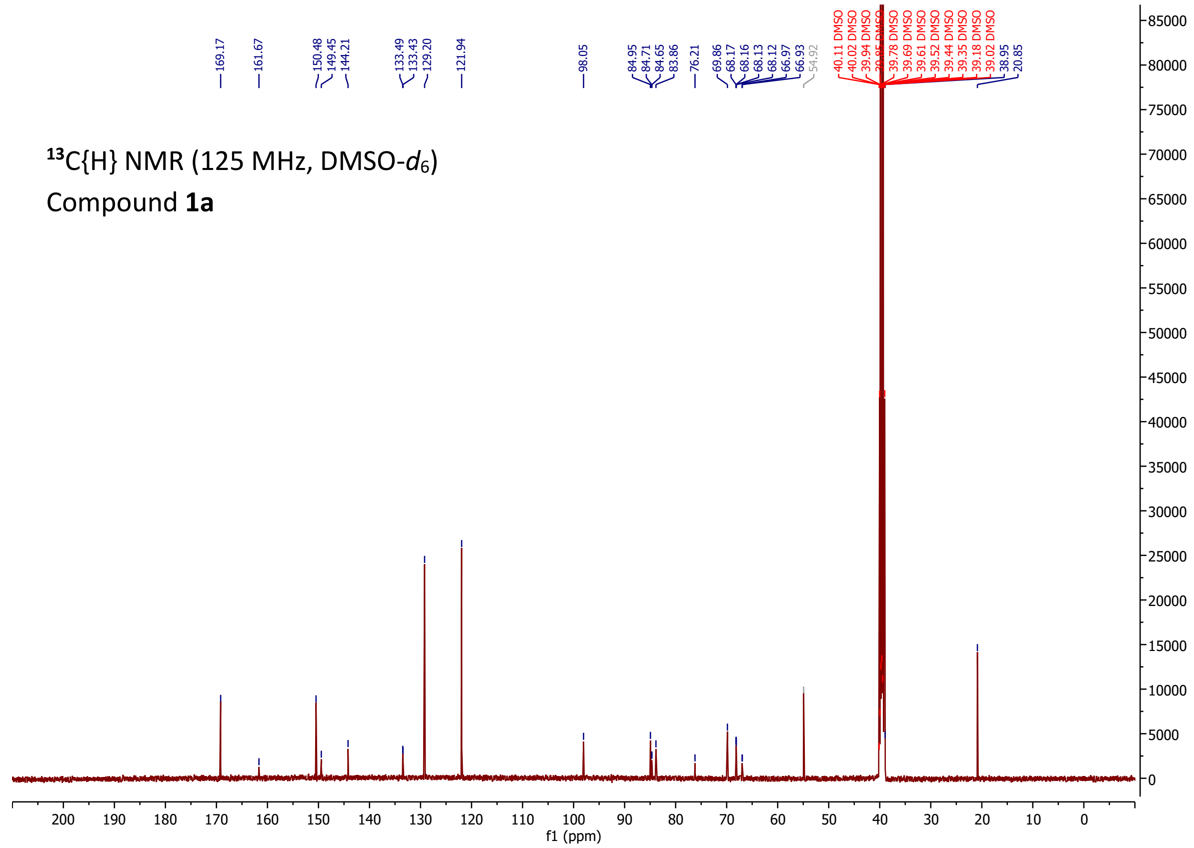

13C{H} NMR (125 MHz, DMSO-d6)
Compound 1a
169.17
161.67
150.48
149.45
144.21
133.49
133.43
129.20
121.94
98.05
84.95
84.71
84.65
83.86
76.21
69.86
68.17
68.16
68.13
68.12
66.97
66.93
54.92
40.11 DMSO
40.02 DMSO
39.94 DMSO
39.78 DMSO
39.69 DMSO
39.61 DMSO
39.52 DMSO
39.44 DMSO
39.35 DMSO
39.18 DMSO
39.02 DMSO
38.95
20.85
f1 (ppm)

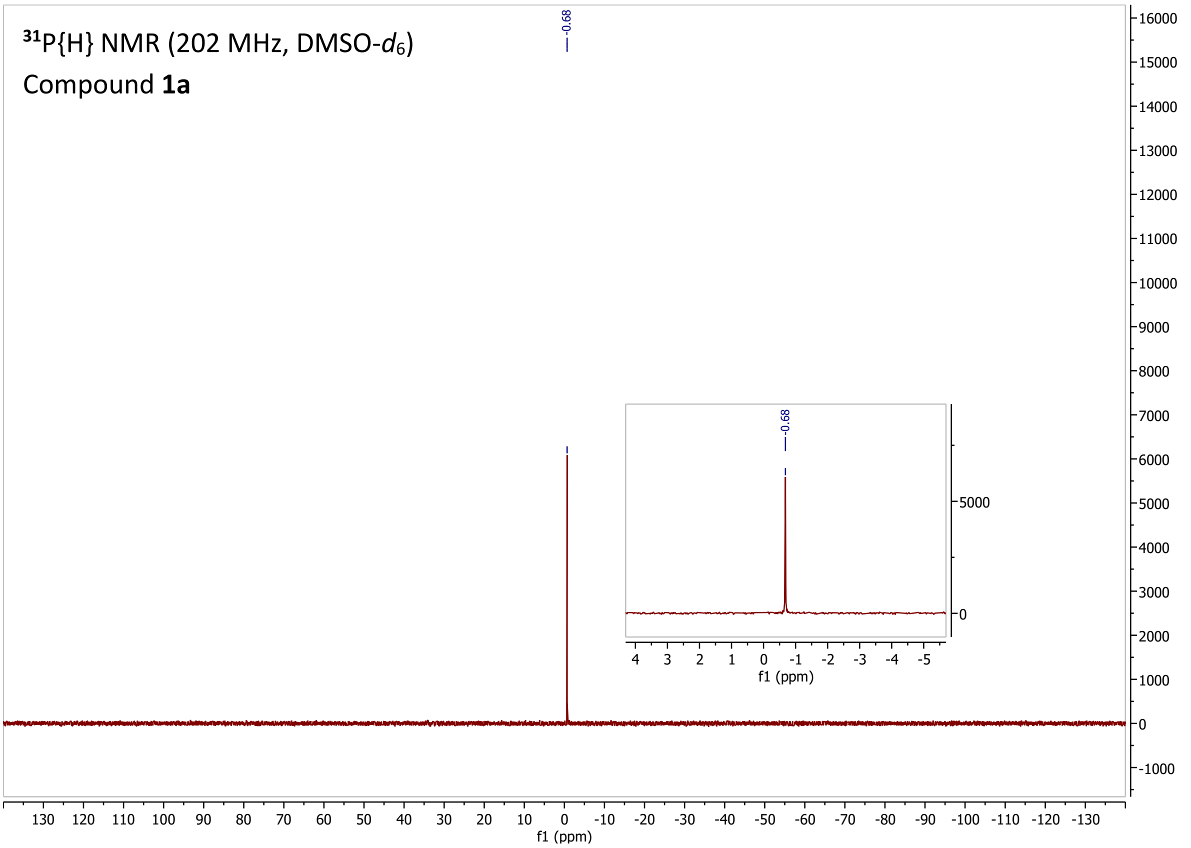

$^{31}P\{H\}$ NMR (202 MHz, DMSO-$d_6$)
Compound **1a**
-0.68
f1 (ppm)

$^{1}H$ NMR (500 MHz, DMSO-$d_6$)

Compound **1b**

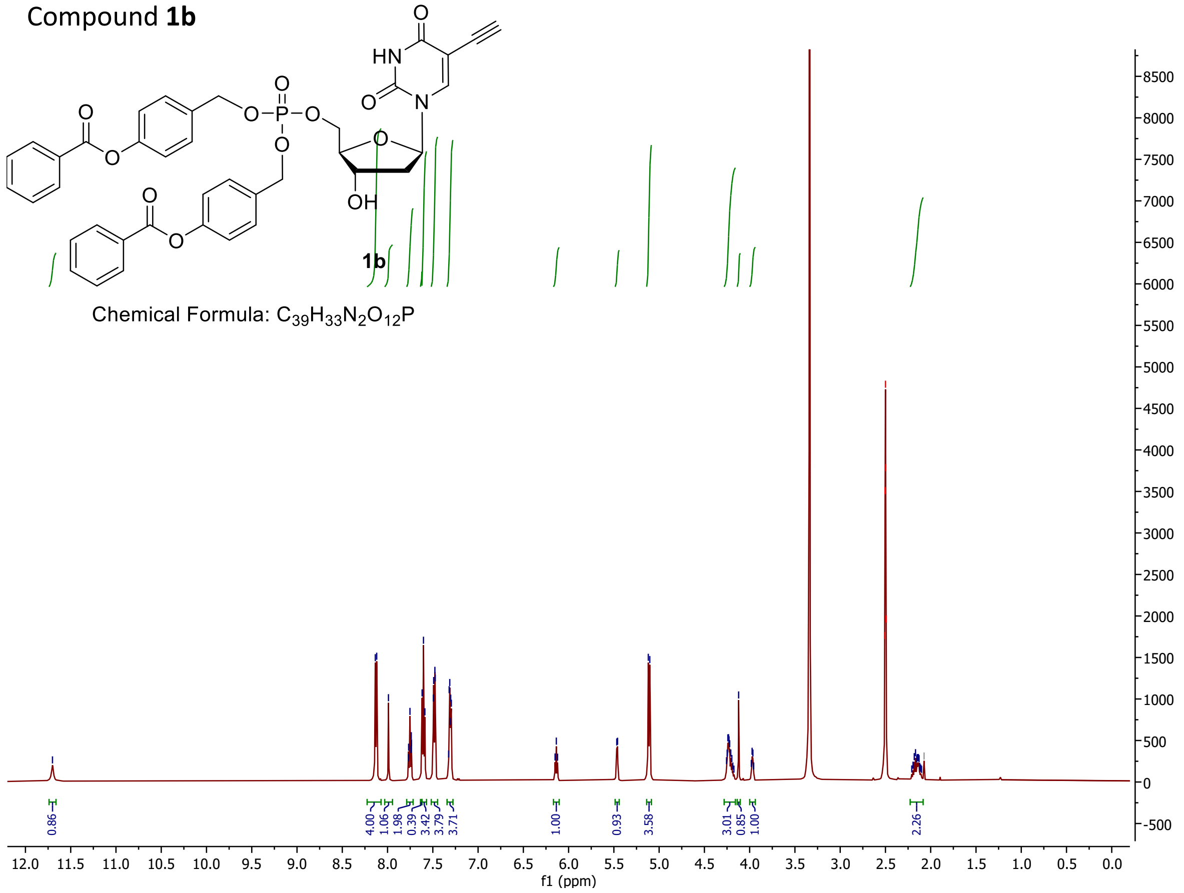


$^{13}C\{H\}$ NMR (125 MHz, DMSO-$d_6$)

Compound **1b**

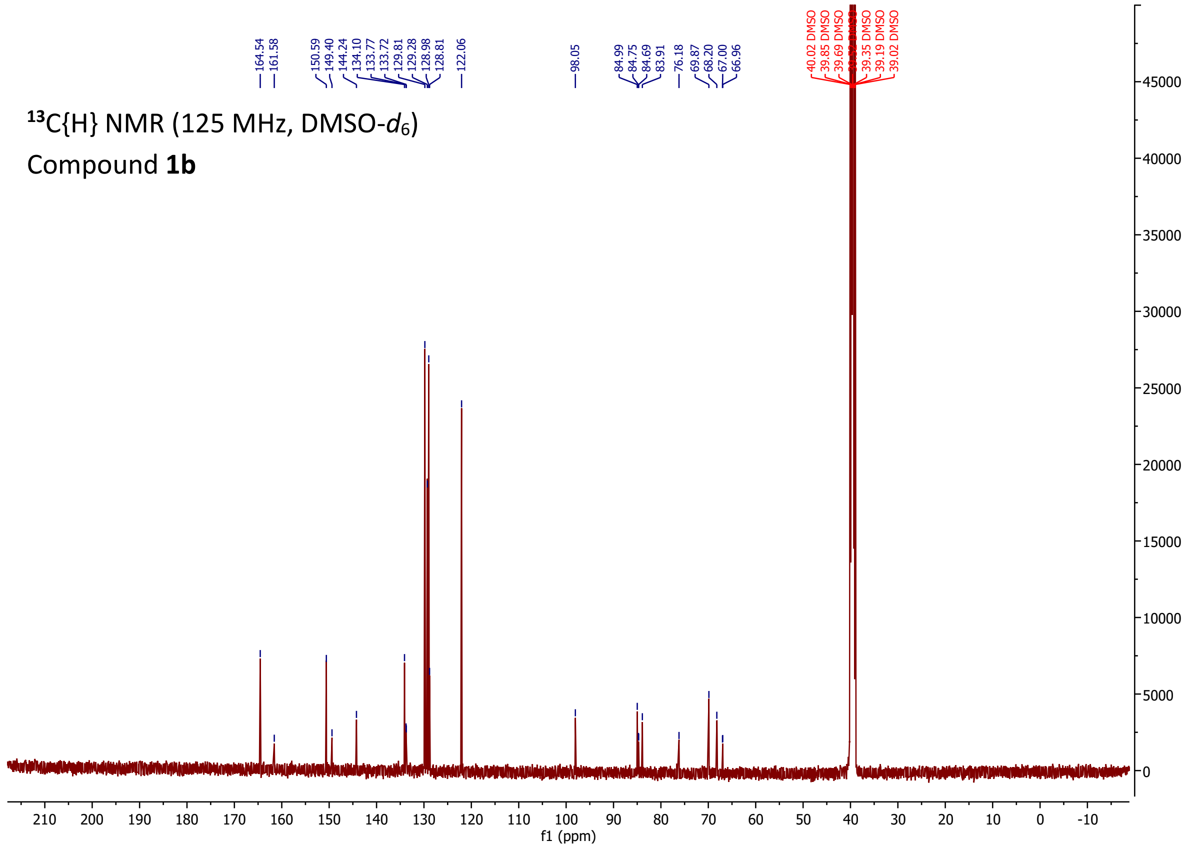

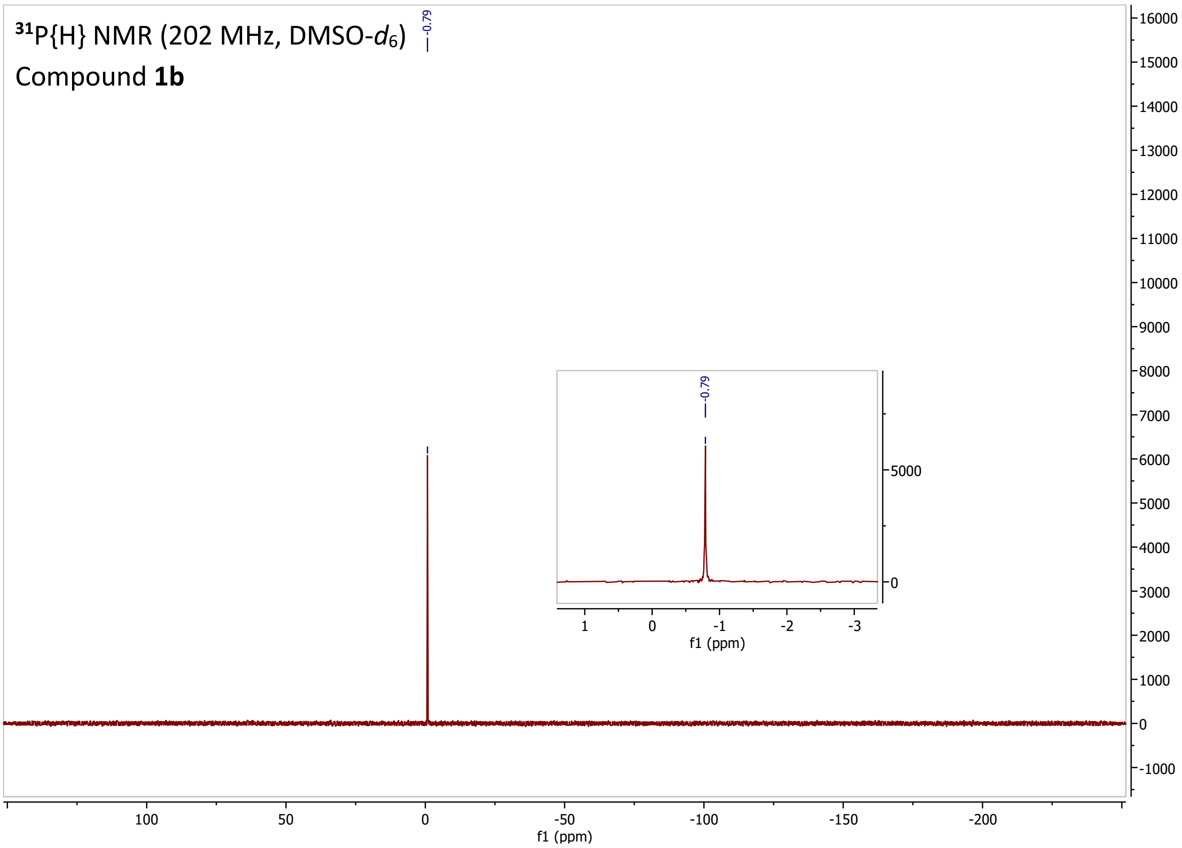

$^{31}$P{H} NMR (202 MHz, DMSO-$d_6$)
Compound **1b**
-0.79
16000
15000
14000
13000
12000
11000
10000
9000
8000
7000
6000
5000
4000
3000
2000
1000
0
-1000
100
50
0
-50
-100
-150
-200
f1 (ppm)
-0.79
5000
0
1
0
-1
-2
-3
f1 (ppm)

$^{1}H$ NMR (500 MHz, DMSO-$d_6$)

Compound **1c**

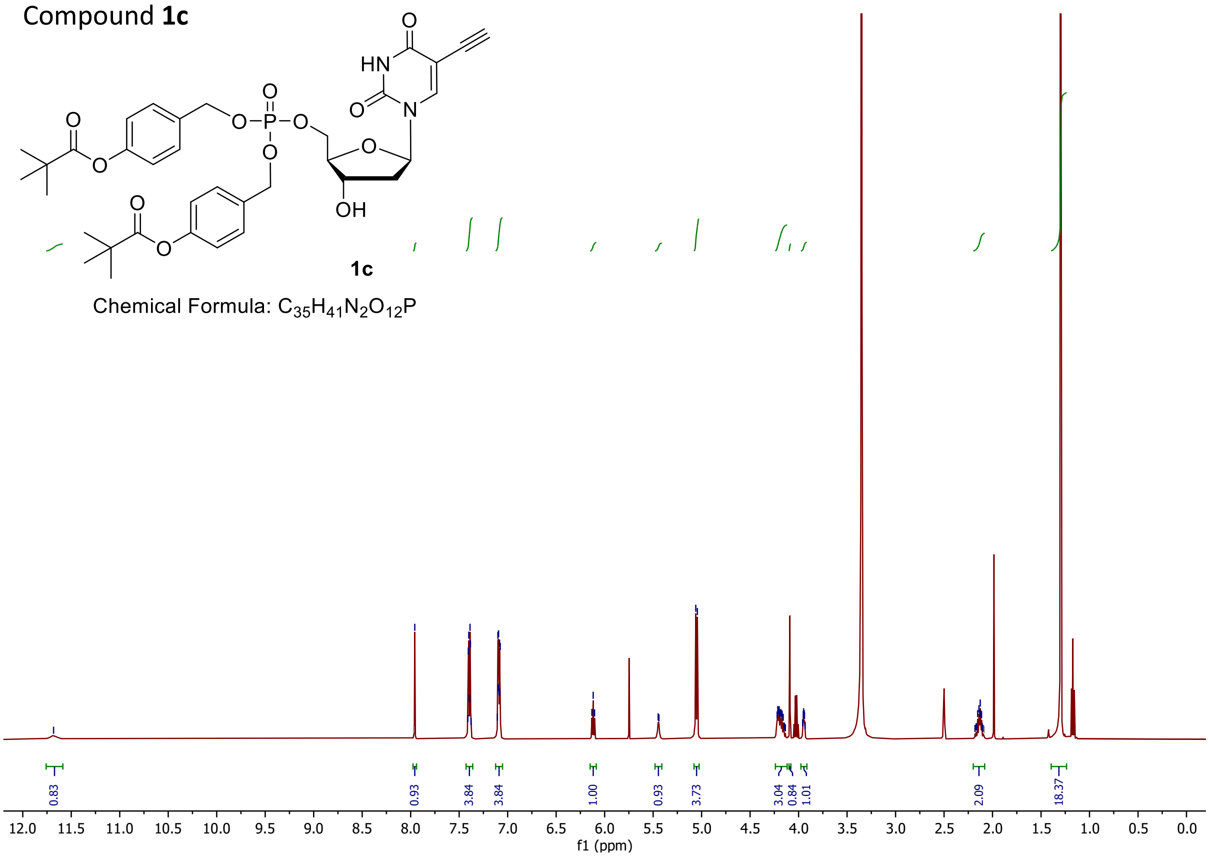


$^{13}C\{H\}$ NMR (125 MHz, DMSO-$d_6$)

Compound **1c**

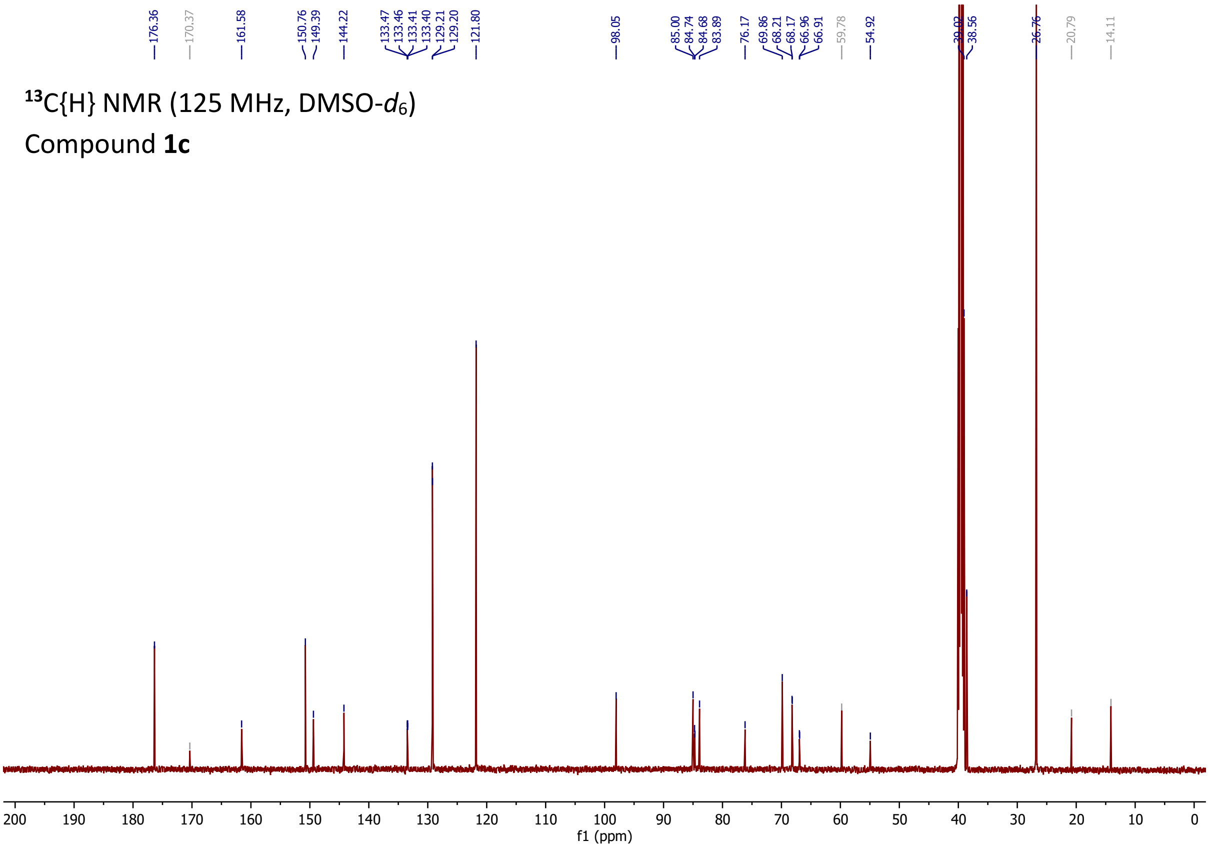

$^{31}P\{H\}$ NMR (202 MHz, DMSO-$d_6$)

Compound **1c**

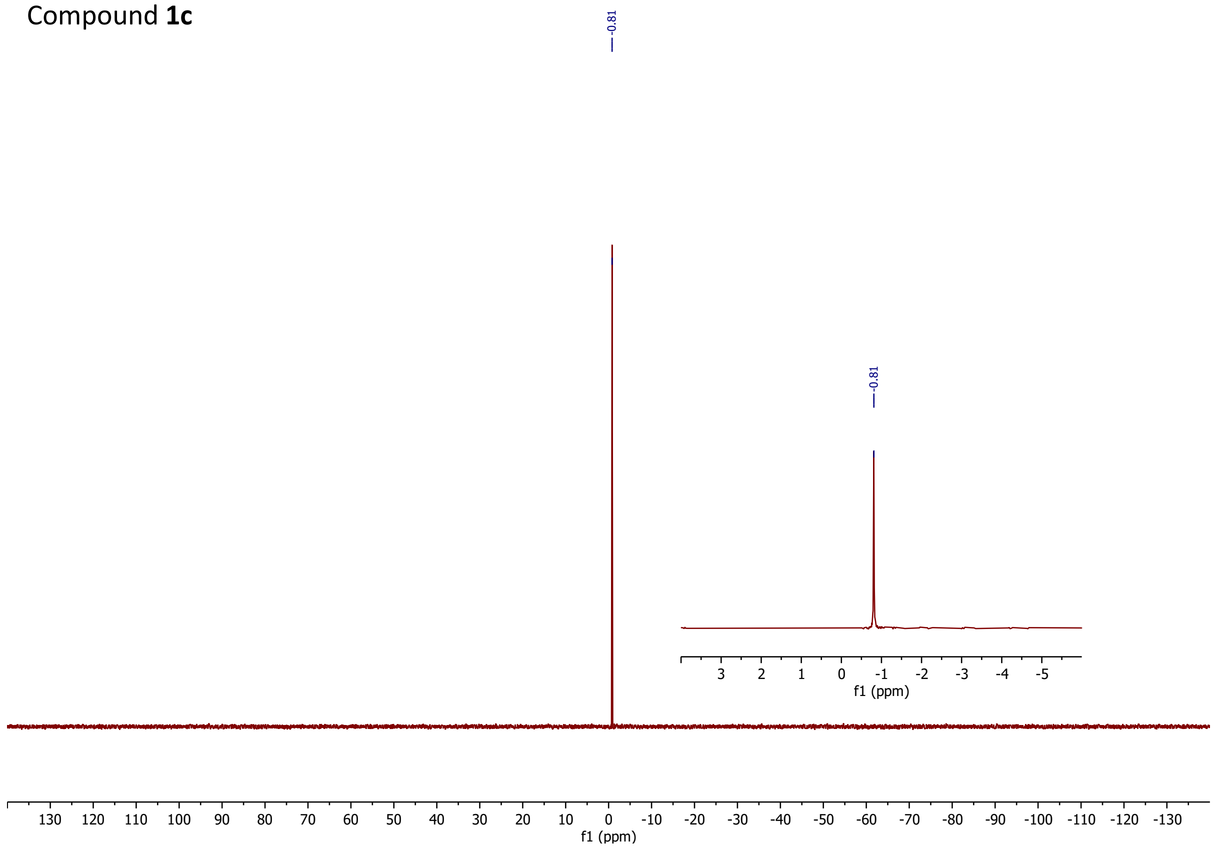

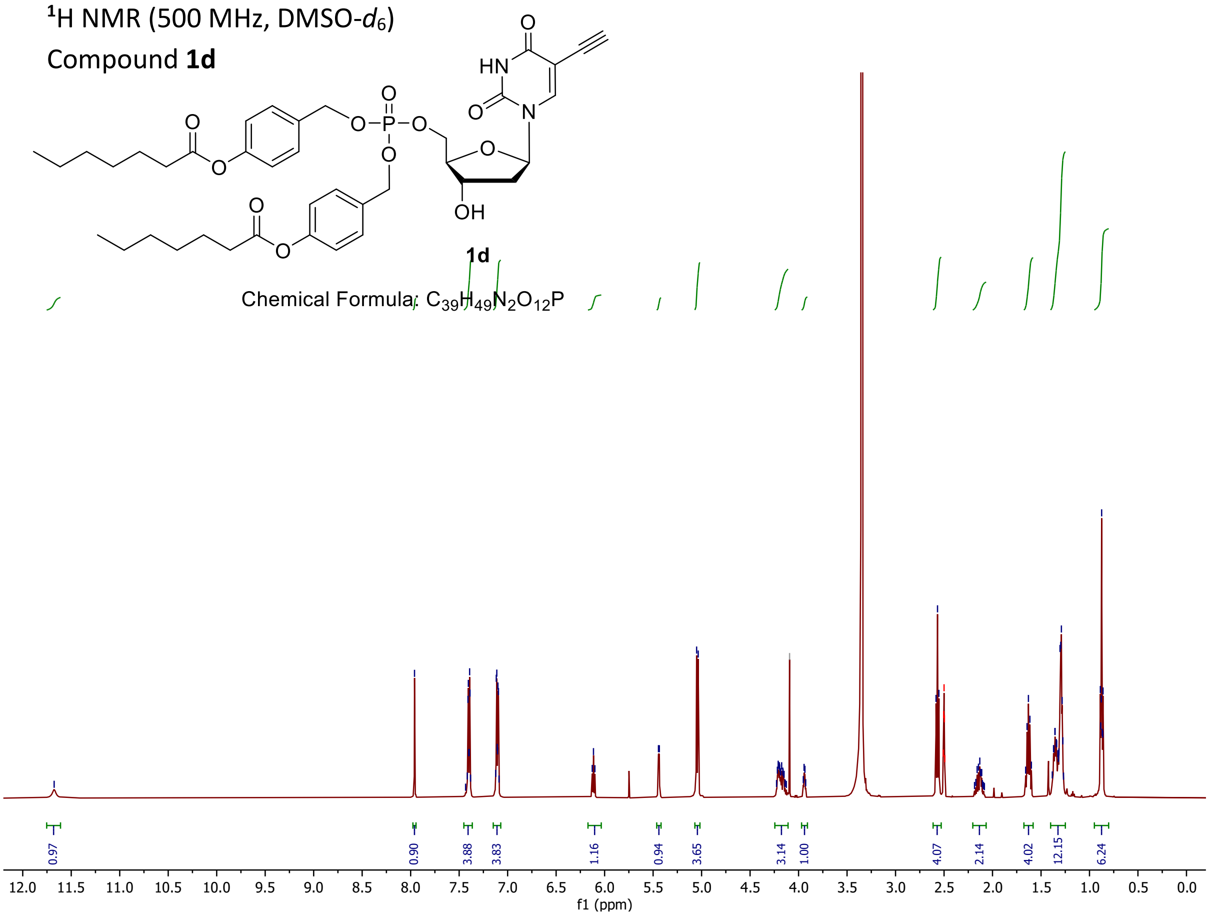

¹H NMR (500 MHz, DMSO-d6)
Compound 1d
1d
Chemical Formula: C39H49N2O12P
0.97
0.90
3.88
3.83
1.16
0.94
3.65
3.14
1.00
4.07
2.14
4.02
12.15
6.24
f1 (ppm)


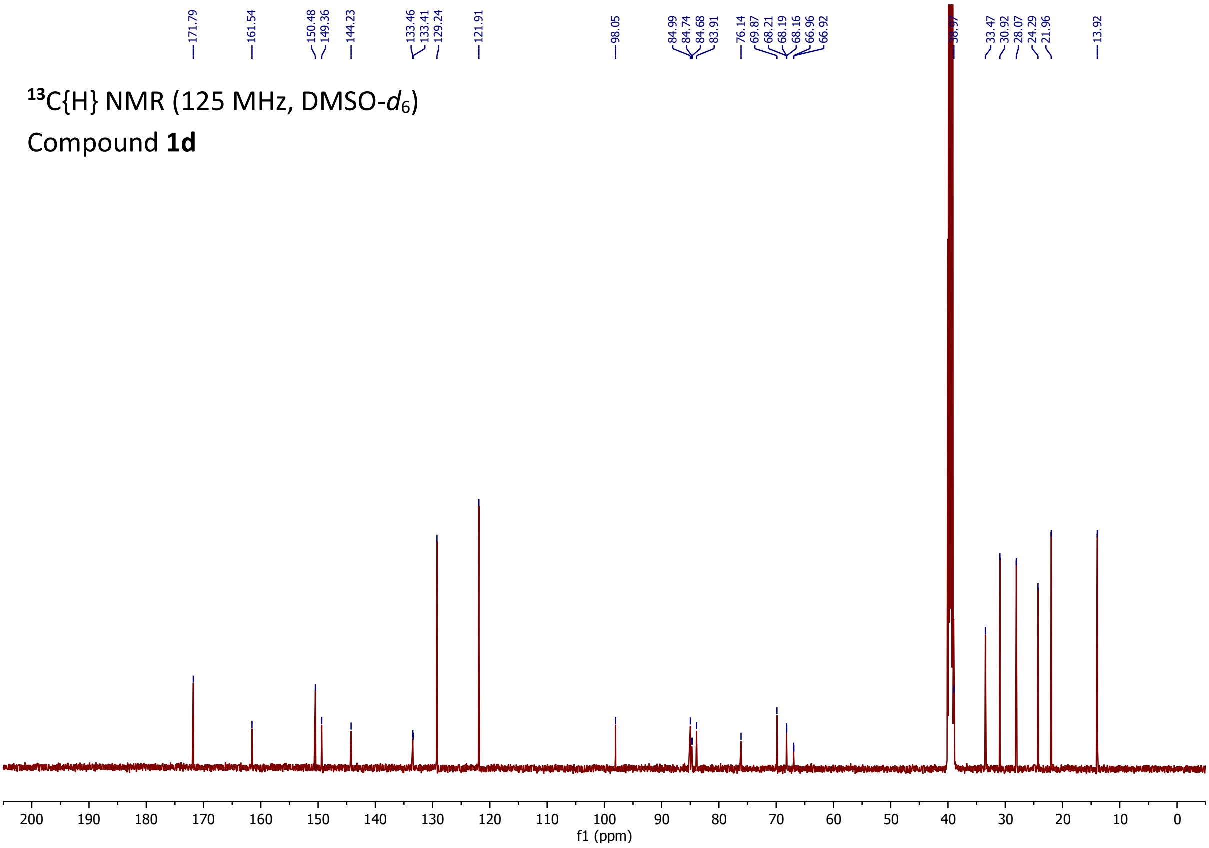

¹³C{H} NMR (125 MHz, DMSO-d6)
Compound 1d
171.79
161.54
150.48
149.36
144.23
133.46
133.41
129.24
121.91
98.05
84.99
84.74
84.68
83.91
76.14
69.87
68.21
68.19
68.16
66.96
66.92
33.47
30.92
28.07
24.29
21.96
13.92
f1 (ppm)

$^{31}P\{H\}$ NMR (202 MHz, DMSO-$d_6$)
Compound **1d**

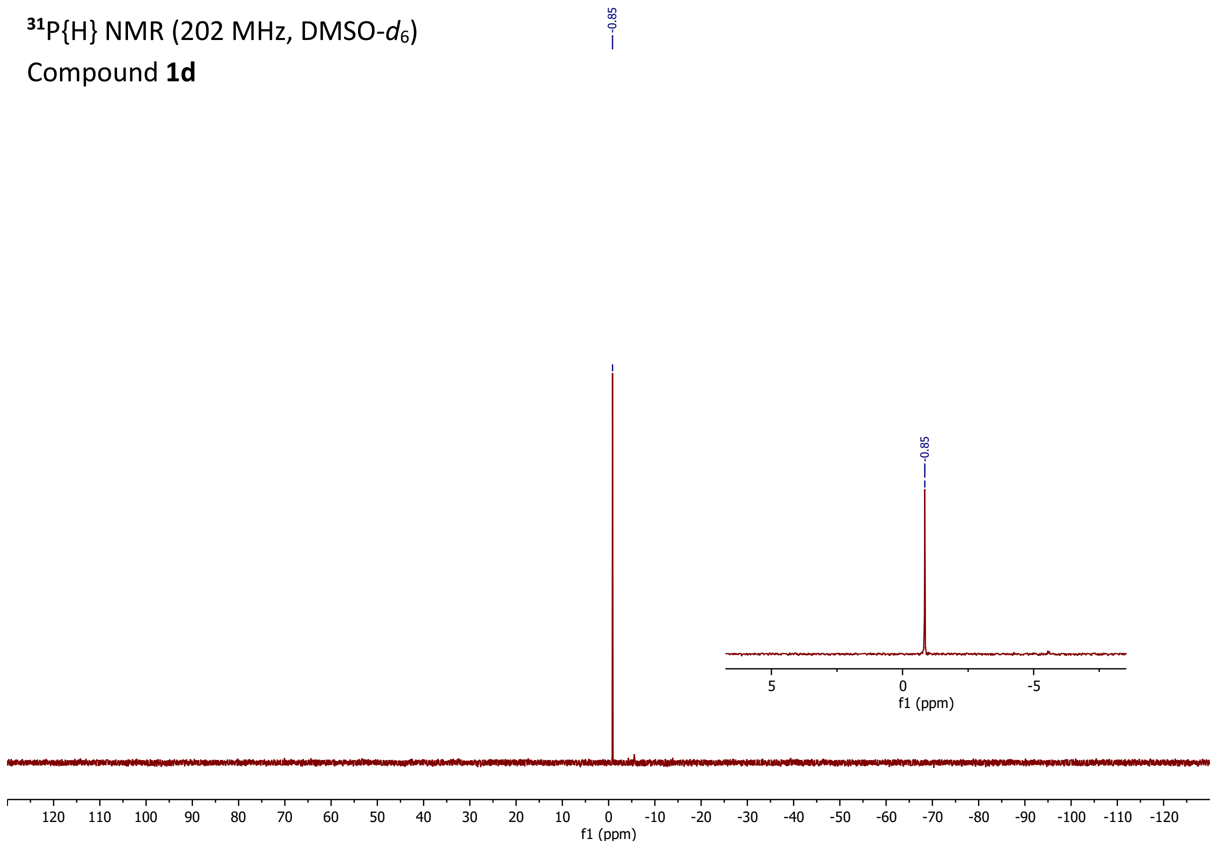

**Media stability studies of chemical probes 1a-1d**

**Methods:** Stock solutions of compounds **1a–1d** (10 mM) were prepared in 50 % (*v/v*) acetonitrile/methanol (both HPLC grade) and stored in glass vials with PTFE-lined caps at 4 °C. Working solutions (1.0 mM) were prepared by diluting the stocks in 100% acetonitrile and were stored similarly. Analysis was performed using LC-MS/MS (Waters 2795 solvent delivery system linked to a MicroMass Quattro Micro tandem MS/MS), conducted in positive ion multiple reaction monitoring mode (MRM) with argon as collision gas at approx. $5 \times 10^{-3}$ mbar. Optimum LC-MS/MS conditions were established for each compound by direct infusion of solutions of each compound (approx. 2.0 µg/mL; 2.5 – 3.3 µM) prepared in 50 % (*v/v*) acetonitrile/$H_2O$ containing 0.1 % (*v/v*) formic acid and 100 µM sodium acetate) into the LC-MS/MS. For the four compounds, capillary voltages ranged from 2.65 – 3.00 kV, cone voltages were 27 – 40 V and collision energies between 21 and 32 eV. Source and desolvation temperatures were set at 140 °C and 350 °C, respectively, with the desolvation gas (nitrogen) set to 400 L/h. Parent ions for all four compounds were measured as their respective sodium adducts. To ensure adequate sensitivity of analysis, multiple mass transitions from the parent adducts were summed for all compounds, Table S1.

Chromatography was conducted using a $C_{18}$ column (Luna, 50 × 4.6 mm × 3.0 µm, Phenomenex) maintained at 37 °C, with a mobile phase flow rate of 0.7 mL/min. For compounds **1a**, **1b** and **1c**, isocratic elution with an acetonitrile/$H_2O$ mobile phase (70 % acetonitrile : 30 % $H_2O$ (*v/v*) containing 0.1 % formic acid and 100 µM sodium acetate (to assist with sodium adduct formation)) was used. For compound **1d**, a gradient elution between 60 % and 95 % acetonitrile over 3.5 min was employed. Under these conditions, retention times ranged between 1.13 – 2.91 min for compounds **1a-1d**, and 1.17 min for the internal standard.

Stability studies were conducted in triplicate using RPMI-complete media (RPMI with sodium bicarbonate, 25 mM HEPES, 50 µg/mL hypoxanthine, 5% human serum and 2.5 mg/mL albumax II) in 50 mL polypropylene conical-bottom tubes, maintained at 37 °C in an oscillating water bath (Grant OLS26; 60 RPM). The media (total volume = 3.0 mL) was pre-warmed at 37 °C for 10 min before being spiked with 15 µL of the 1.0 mM solutions described above to provide a starting concentration of 5.0 µM. Final organic (acetonitrile) concentration in the incubation tubes was 0.5 %. Samples (100 µL) were removed at t = 0 (immediately after reaction start) and then at selected times at half-hourly or hourly intervals (for compounds **1b-**

**1d**; more rapidly up to 1 h for compound **1a**), up to 6 h post-initiation and were added to microcentrifuge tubes containing 300 µL of ice-cold acetonitrile containing internal standard (clotrimazole (Sigma Aldrich, Sydney, Australia), 50 ng/mL + 0.1 % (*v/v*) formic acid + 100 µM sodium acetate). The tubes were vortex mixed for 30 s and centrifuged (13000 × *g*, 5 min, rt) and ~180 µL of the supernatant transferred to polypropylene 96-well plates for analysis.

Peak area ratios (compounds **1a-1d**: internal standard) were determined at each timepoint for each tube and expressed as percent remaining (normalised to t = 0 as 100%; Table S2). Percent remaining *versus* time data were fitted to an exponential decay function to determine the first-order rate constant (*k*) for substrate depletion. The rate of depletion for each sample was used to calculate the half-life of degradation according to the equation:

$$t_{½} = \frac{\ln(2)}{k}$$

To ensure adequate linearity of the LC-MS/MS analysis of all compounds over an appropriate range, standard curves were prepared in RPMI-complete media on ice at 0.5, 1.0, 2.0, 3.0, 4.0, 5.0 and 7.5 µM and analysed prior to any stability studies being conducted. Linearity of analysis for all compounds over that range was confirmed with $r^2$ of > 0.99 in each case. The lower concentration of 0.5 µM represents 10 % of the starting concentration of the stability studies.

**Table S1:** Mass transitions summed for compounds **1a-1d**.

| **Compound** | **1a** | **1b** | **1c** | **1d** |
|---|---|---|---|---|
| **Mass Transitions** | *m/z* 651.1>348.9 | *m/z* 775.1>410.5<br>*m/z* 775.1>532.4<br>*m/z* 775.1>639.4 | *m/z* 735.2>386.5<br>*m/z* 735.2>391.1<br>*m/z* 735.2>599.1 | *m/z* 790.8>354.9<br>*m/z* 790.8>419.1<br>*m/z* 790.8>654.9 |

**Table S2:** Degradation of compounds **1a-1d** in RPMI-complete media at 37 °C.

| **Time (h)** | **% Remaining[1]** | | | | | | | |
|---|---|---|---|---|---|---|---|---|
| | **Compound 1a** | | **Compound 1b** | | **Compound 1c** | | **Compound 1d** | |
| | **Mean[2]** | **SD[2]** | **Mean** | **SD** | **Mean** | **SD** | **Mean** | **SD** |
| 0 | 100 | 0 | 100 | 0 | 100 | 0 | 100 | 0 |
| 0.167 | 22.9 | 0.81 | --- | --- | --- | --- | --- | --- |
| 0.333 | 5.87 | 0.43 | --- | --- | --- | --- | --- | --- |
| 0.5 | 1.70 | 0.31 | 98.0 | 14.2 | --- | --- | --- | --- |
| 1 | BLQ[3] | BLQ | 83.5 | 6.33 | 72.7 | 8.91 | 29.6 | 0.49 |
| 2 | ---[4] | --- | 55.9 | 5.90 | 50.3 | 2.45 | 7.37 | 1.21 |
| 3 | --- | --- | --- | --- | 32.8 | 1.67 | 2.13 | 0.46 |
| 4 | --- | --- | 22.3 | 1.21 | 18.8 | 0.11 | BLQ | BLQ |
| 6 | --- | --- | 8.09 | 0.733 | 12.2 | 2.11 | --- | --- |

[1]Normalized to t = 0. [2] Mean ± SD (n=3). [3]Below lower limit of quantitation. [4]No sample taken at this timepoint.

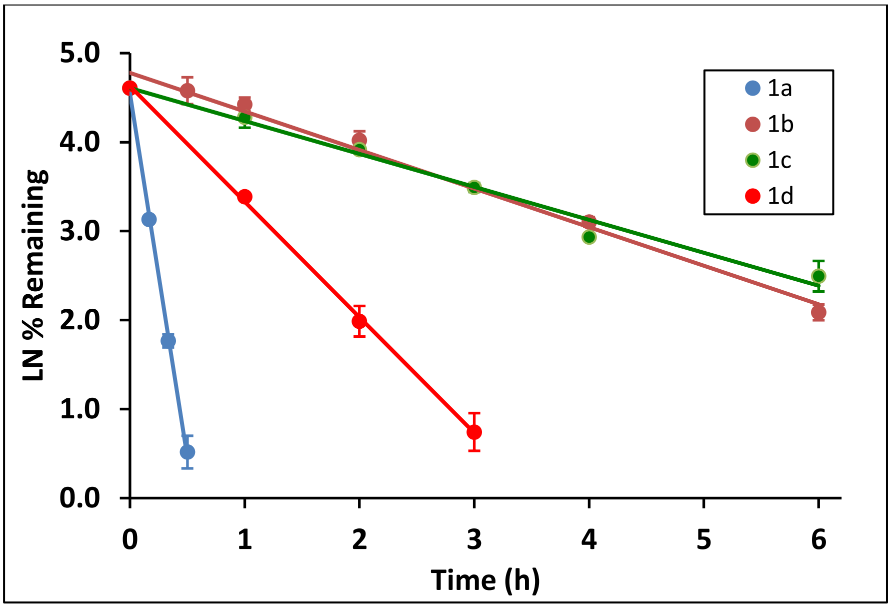


**Figure S1:** Stability of compounds **1a-1d** in RPMI-complete media at 37 °C. LN-transformed data; mean ± SD (n = 3).

**1a**: Y = -8.1743x + 4.5465; $R^2$ = 0.9985; t½ = 0.085 ± 0.0004 h.

**1b**: Y = 0.4333x + 4.778; $R^2$ = 0.9883; t½ = 1.60 ± 0.05 h.

**1c**: Y = -0.3702x + 4.6076; $R^2$ = 0.9837; t½ = 1.88 ± 0.14 h.

**1d**: Y = -1.2988x + 4.6291; $R^2$ = 0.9993; t½ = 0.53 ± 0.03 h.
